# Hepatocyte FAM210A deficiency disrupts mitochondrial function and triggers juvenile steatosis with compensatory repair in adulthood

**DOI:** 10.1101/2025.09.02.673111

**Authors:** Yubo Wang, Leah Chu, Yumei Zhou, Zihao Jin, Kyoungrae Kim, Katie Heiden, Miguel A. Gutierrez-Monreal, Junxiao Ren, Yufen Li, Zhiyong Cheng, Kari B. Basso, Karyn A. Esser, Terence E. Ryan, Feng Yue

## Abstract

Mitochondrial dynamics are central to maintaining liver metabolic homeostasis, yet the mechanisms that safeguarding mitochondrial integrity during development and metabolic dysfunction remain poorly defined. Here, we identify Family with sequence similarity 210 member A (FAM210A) as a hepatocyte-enriched mitochondrial regulator essential for postnatal liver maturation. Hepatocyte-specific deletion of *Fam210a* (*Fam210a^HKO^*) in mice caused early growth restriction, reduced body and liver mass, and pronounced hepatic steatosis with glycogen depletion. These defects were accompanied by lower postprandial glucose levels in the fasted–refeeding state, impaired oxidative phosphorylation, reduced mtDNA content, and abnormal cristae architecture. Transcriptomic and proteomic profiling revealed broad suppression of fatty acid, sterol, and bile acid metabolism, with concomitant glutathione stress responses. Mechanistically, FAM210A deficiency disrupted the YME1L–OPA1 axis, driving excessive OPA1 cleavage and cristae destabilization. Strikingly, these juvenile defects were transient and resolved by adulthood, underpinned by enhanced hepatocyte proliferation and mitochondrial biogenesis, consistent with a compensatory stress-adaptive response via the activation of ISR signaling. Together, these findings uncover FAM210A as a developmental safeguard of mitochondrial remodeling in hepatocytes and indicate compensatory programs with therapeutic relevance for chronic liver disease.

## Introduction

The liver is a central metabolic organ responsible for coordinating daily and lifelong nutrient utilization, detoxification, and systemic energy balance^1,2^. Whereas the fetal liver serves as the principal site for hematopoiesis, hepatocytes undergo rapid and profound metabolic maturation during postnatal development, acquiring the capacity to primarily regulate lipid, glucose, and amino acid metabolism while maintaining redox and immune homeostasis^3–5^. Given its continuous exposure to dietary fluctuations, xenobiotics, and metabolic by-products, the liver is highly susceptible to stress and injury^6,7^. However, it has evolved a remarkable regenerative capacity, engaging proliferative and adaptive programs that restore tissue mass and function following toxin exposure or metabolic overload^8,9^. Understanding the molecular factors that couple metabolic homeostasis to hepatic injury and repair is therefore critical for advancing liver biology and developing new therapeutics for liver disease.

Mitochondria are indispensable for liver physiology, fueling oxidative phosphorylation, fatty acid oxidation, gluconeogenesis, and urea cycle activity while also controlling redox balance and apoptotic signaling^10,11^. The structural plasticity of mitochondria, through cristae remodeling, biogenesis, mitophagy, and dynamic fusion–fission cycles, establishes its energy metabolic capacity and enables hepatocytes to adapt to changing nutritional and physiological demands during postnatal development and injury^12,13^. Disruption of mitochondrial homeostasis is a hallmark of diverse hepatic disorders, including steatosis, insulin resistance, and liver failure^11,14,15^. Conversely, mitochondrial stress responses, including activation of the integrated stress response (ISR) and mitochondrial unfolded protein response (UPRmt), are increasingly recognized as key drivers of hepatic adaptation and regeneration^16–20^. Thus, mitochondria act not only as bioenergetic powerhouses but also as signaling platforms that integrate metabolic cues with injury and repair processes in the liver.

Family with sequence similarity 210 member A (FAM210A) has emerged as a novel mitochondrial regulator with essential implications for metabolism, organ function, and systemic physiology^21^. Initially identified by genome-wide association studies linking genetic variation near FAM210A to bone mineral density and lean mass in humans^22–24^, recent genetic studies from our and other groups cross different species demonstrate its critical roles in the heart^25,26^, skeletal muscle^27,28^, adipose tissue^29,30^, and reproductive organs^31^. Loss of FAM210A causes impaired brown fat thermogenesis and cold intolerance, dilated cardiomyopathy, and progressive myopathy with reduced strength^21^. At the cellular level, FAM210A deficiency leads to fragmented mitochondria, disrupted cristae, reduced mitochondrial content, and impaired respiration in myofibers, adipocytes, and cardiomyocytes, ultimately compromising energy homeostasis^25–30^. Mechanistically, FAM210A acts in a tissue-dependent manner: in brown adipocytes, it regulates OPA1 processing through the YME1L/OMA1 axis to control cristae remodeling^29^; in cardiomyocytes, it interacts with EF-Tu to support mitochondrial translation and contractile function^25,26^; and in skeletal muscle, it modulates cytosolic protein synthesis via acetyl-CoA metabolism and ribosomal activity^28^. Collectively, FAM210A functions as a multifaceted hub integrating mitochondrial proteostasis, inter-organelle crosstalk, and metabolism. Despite growing recognition of its importance in maintaining mitochondrial integrity, the role of FAM210A in the liver, an organ uniquely relying on mitochondrial function during development, homeostasis and injury, remains unexplored.

In this study, we investigated the role of FAM210A in the liver by generating a hepatocyte-specific *Fam210a* knockout (*Fam210a^HKO^*) mouse model and tracking its impact across postnatal development. Loss of *Fam210a* impedes liver growth and impairs hepatic metabolism and mitochondrial function during the juvenile stage. By integrating transcriptomic and mitochondrial proteomics analyses with biochemical and ultrastructural analyses, we explored the molecular mechanisms underlying the defects of liver metabolism upon *Fam210a* deletion with the focus on the regulation of mitochondrial remodeling. By assessing liver homeostasis in FAM210A-deficient mice over time, we further identified compensatory programs that restore metabolic homeostasis in adulthood. Our findings reveal a developmental window during which mitochondrial integrity is particularly critical and establish a framework for understanding how stress-adaptive pathways preserve liver function in the absence of FAM210A.

## Results

### *Fam210a* is essential for postnatal liver development

We first evaluated the expression pattern of FAM210A in the liver across postnatal developmental stages. The protein level of FAM210A was markedly elevated from early postnatal stages to adulthood (Fig. 1a), indicating its progressive upregulation during liver maturation. This expression pattern is consistent with the increasing metabolic demands and mitochondrial activity during hepatic development^3,32^, highlighting a potential role for FAM210A in the functional maturation of the liver. To determine the cell populations expressing *Fam210a* in the heterogenous liver tissue^33,34^, we reanalyzed the published single-cell transcriptomics (GSE109774)^35^ and found that *Fam210a* expression is predominantly confined to hepatocytes, with minimal signal in non-parenchymal cell types (Fig. 1b).

**Fig.1.**
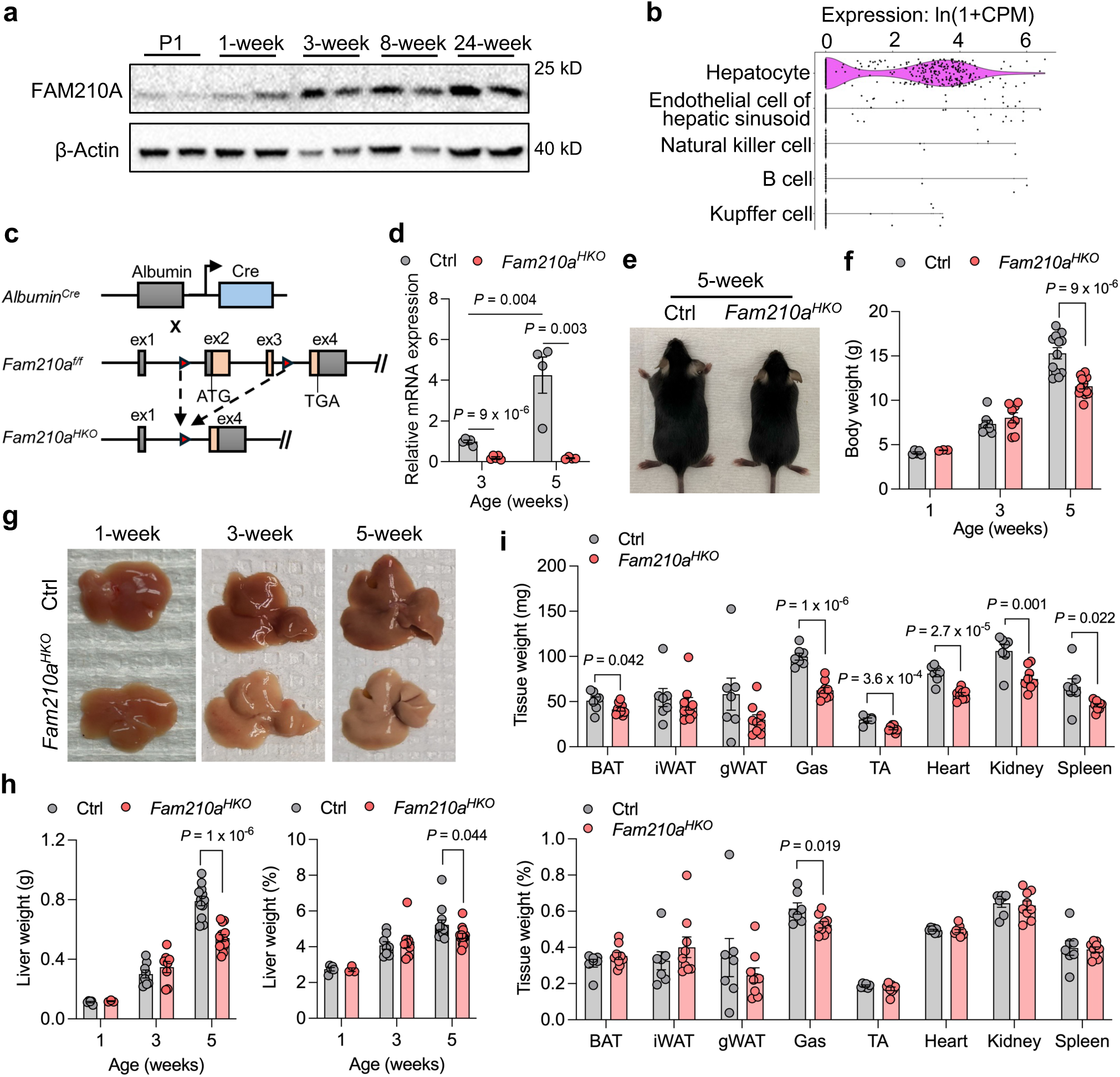
Hepatocyte-specific deletion of *Fam210a* impedes juvenile growth and liver development. **a** Immunoblot showing FAM210A protein levels in mouse liver at different ages. **b** Single-cell RNA-seq analysis indicating predominant expression of *Fam210a* mRNA in hepatocytes. **c** Schematic illustration of the strategy used to generate hepatocyte-specific *Fam210a* knockout (*Fam210a^HKO^*) mice. ex, exon. **d** Relative hepatic *Fam210a* mRNA expression in Ctrl and *Fam210a^HKO^* mice at 3 and 5 weeks old (n = 4; mean ± s.e.m.; two-tailed, unpaired Student’s t-test). **e** Representative images of Ctrl and *Fam210a^HKO^* mice at 5 weeks old. **f** Body weight of Ctrl and *Fam210a^HKO^* mice at 1, 3, and 5 weeks old (1-week-old: Ctrl, n = 5; *Fam210a^HKO^*, n = 3; 3-week-old: Ctrl and *Fam210a^HKO^*, n = 8; 5-week-old: Ctrl, n = 12; *Fam210a^HKO^*, n = 14; mean ± s.e.m.; two-tailed, unpaired Student’s t-test). **g** Representative liver morphology of Ctrl and *Fam210a^HKO^* mice at 1, 3, and 5 weeks old. **h** Liver weight and liver-to-body weight ratio in Ctrl and *Fam210a^HKO^* mice at different ages (1-week-old: Ctrl, n = 5; *Fam210a^HKO^*, n = 3; 3-week-old: Ctrl and *Fam210a^HKO^*, n = 8; 5-week-old: Ctrl, n = 12; *Fam210a^HKO^*, n = 14; mean ± s.e.m.; two-tailed, unpaired Student’s t-test). **i** Tissue weights and percentages of other major organs in Ctrl and *Fam210a^HKO^* mice at 5 weeks old (Ctrl, n = 7; *Fam210a^HKO^*, n = 9; mean ± s.e.m.; two-tailed, unpaired Student’s t-test).

To understand the physiological function of FAM210A in the liver, we generated *Fam210a^HKO^* mice by crossing *Fam210a^f/f^*mice with *Albumin-Cre* (*Alb-Cre*) transgenic mice (Fig. 1c). Quantitative qPCR confirmed efficient deletion of *Fam210a* in the *Fam210a^HKO^* liver at 3 and 5 weeks old, while the mRNA expression of *Fam210a* increased significantly in the livers of control mice during this postnatal growth (Fig. 1d). *Fam210a^HKO^*mice were born normally and exhibited comparable body weight before 3 weeks of age, but markedly decreased body weight at 5 weeks (Fig. 1e, f). Notably, the livers of *Fam210a^HKO^* mice appeared abnormally pale and significantly smaller at 5 weeks old compared with littermate controls (Fig. 1g, h). The reduction in liver mass was accompanied by decreases in adipose tissue, skeletal muscle, and other organs, including the heart, kidneys, and spleen in *Fam210a^HKO^* mice (Fig. 1i). These results suggest that hepatic FAM210A deficiency impedes juvenile growth and liver development during early postnatal stage.

### *Fam210a* deficiency disrupts hepatic lipid and glucose metabolism in juvenile mice

To determine whether *Fam210a* KO influences liver histology, we performed H&E staining and observed lipid accumulation in liver sections of *Fam210a^HKO^* mice starting at 3 weeks old while remarkably increased at 5 weeks old, when compared to the control livers (Fig. 2a). Consistently, BODIPY staining of liver sections confirmed this observation with larger and more numerous lipid droplets observed in *Fam210a^HKO^* mice than in controls at both 3 and 5 weeks old (Fig. 2b). Further quantification of hepatic triglycerides (TAG) showed a 2.5-fold increase in *Fam210a^HKO^* livers compared to controls (200.0 *vs.* 79.6; Fig. 2c). These observations demonstrate an abnormal hepatic lipid accumulation upon *Fam210a* KO at early postnatal stage.

**Fig. 2.**
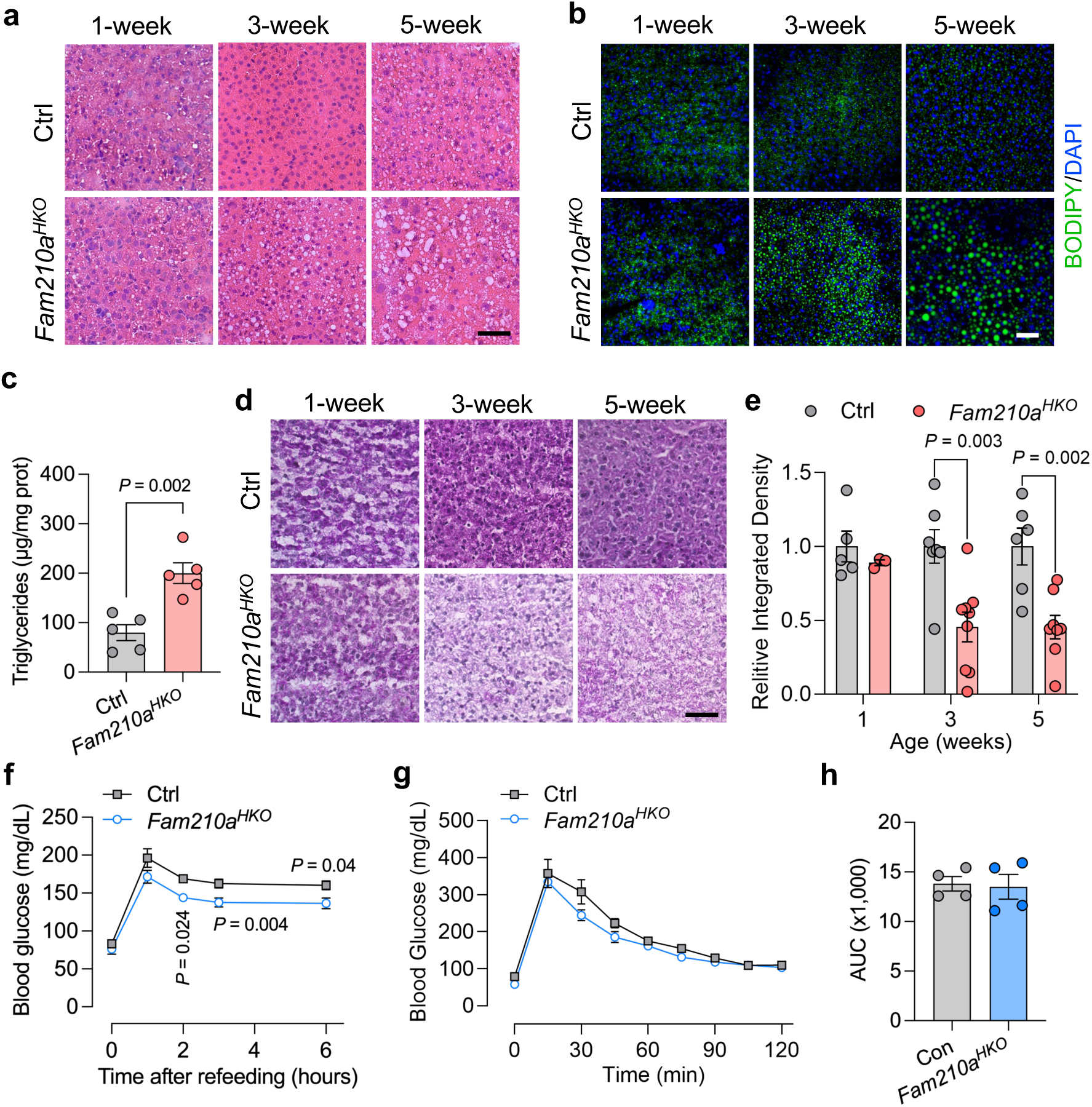
*Fam210a* deficiency causes fat accumulation, low liver glycogen, and hypoglycemia after fasting-refeeding in juveniles. **a** Representative H&E staining of liver sections from Ctrl and *Fam210a^HKO^* mice (scale bar, 50 µm). **b** Representative BODIPY staining of liver sections from Ctrl and *Fam210a^HKO^* mice (scale bar, 50 µm). **c** Hepatic triglyceride levels in Ctrl and *Fam210a^HKO^* mice at 3 weeks old (n = 5; mean ± s.e.m.; two-tailed, unpaired Student’s t-test). **d** Representative PAS staining of liver sections from Ctrl and *Fam210a^HKO^* mice (scale bar, 50 µm). **e** Quantification of relative integrated density in (**d**) (1-week-old: Ctrl, n = 5; *Fam210a^HKO^*, n = 3; 3-week-old: Ctrl, n = 7; *Fam210a^HKO^*, n = 9; 5-week-old: Ctrl, n = 6; *Fam210a^HKO^*, n = 8; mean ± s.e.m.; two-tailed, unpaired Student’s t-test). **f** Blood glucose levels in 5-week-old mice after fast-refeeding (n = 4; mean ± s.e.m.; two-tailed unpaired Student’s t-test). **g** GTT analysis of 5-week-old mice (n = 4; mean ± s.e.m.; two-tailed, unpaired Student’s t-test). **h** Quantification of area under curve (AUC) of GTT analysis in **g** (n = 4; mean ± s.e.m.; two-tailed, unpaired Student’s t-test).

Given the central role of the liver in coordinating lipid and glucose metabolism, we next examined whether hepatic glycogen storage was also affected in *Fam210a^HKO^* mice. Periodic acid–Schiff (PAS) staining revealed that *Fam210a^HKO^* livers appeared clearly paler than controls, indicative of a substantial reduction in hepatic glycogen content (Fig. 2d, e). Liver glycogen metabolism is affected by starvation and refeeding, and the fasting–refeeding cycle serves as a robust physiological test for identifying defects in glycogen metabolism and glucose homeostasis^36^. We performed a fasting–refeeding experiment to further evaluate systemic glucose homeostasis in 5-week-old mice. Following an overnight fast, the blood glucose levels were comparable between the control and *Fam210a^HKO^* mice (Fig. 2f), suggesting that glycogen breakdown and gluconeogenic capacity of *Fam210a*-deficient liver are largely intact during fasting. Intriguingly, while both control and *Fam210a^HKO^* mice reached a peak glucose level at the first hour of refeeding then decreased slowly, the blood glucose level in *Fam210a^HKO^* mice was significantly lower than the control at 2–4 h post-refeeding (Fig. 2f), suggesting that either glucose clearance was enhanced, or hepatic glucose production was reduced. In contrast, intraperitoneal glucose tolerance test (GTT) showed indistinguishable glucose clearance capacity (Fig. 2g-h). Collectively, these results indicate that *Fam210a* KO leads to defective hepatic glycogen storage and impaired postprandial glucose homeostasis.

### Loss of *Fam210a* disrupts hepatic metabolic and immune homeostasis during early postnatal development

To further explore the role of *Fam210a* in liver physiology, we performed transcriptomic profiling of livers from 5-week-old control and *Fam210a^HKO^* mice. Principal component analysis (PCA) revealed clear separation between the two groups, indicating robust transcriptomic divergence (Fig. 3a). Differential expression analysis (log_2_fold change ≥ 1, *Padj* < 0.05) identified 1,023 upregulated and 970 downregulated genes in *Fam210a^HKO^* livers compared to controls (Fig. 3b; Source data 1). KEGG pathway analysis of upregulated genes revealed significant enrichment in pathways related to cell cycle, glutathione metabolism, and p53 signaling pathway, pointing to a stress-adaptive program with activation of glutathione-based redox buffering and repair-linked cell cycle reprogramming (Fig. 3c). Specifically, core antioxidant genes, including glutathione S-transferases (*Gsta2*, *Gsta1*, *Gstm1–4*), glutathione peroxidases (*Gpx3*, *Gpx7*), and microsomal GSTs (*Mgst3*, *Gstp3*) were significantly upregulated in *Fam210a^HKO^* livers (Supplementary Fig. 1a). In contrast, KEGG pathway analysis of downregulated genes revealed significant enrichment in pathways related to complement, sterol metabolism, cytochrome P450–mediated processes, bile acid metabolism, and fatty-acid metabolism (Fig. 3c), indicating broad hepatic metabolic disruption.

**Fig. 3.**
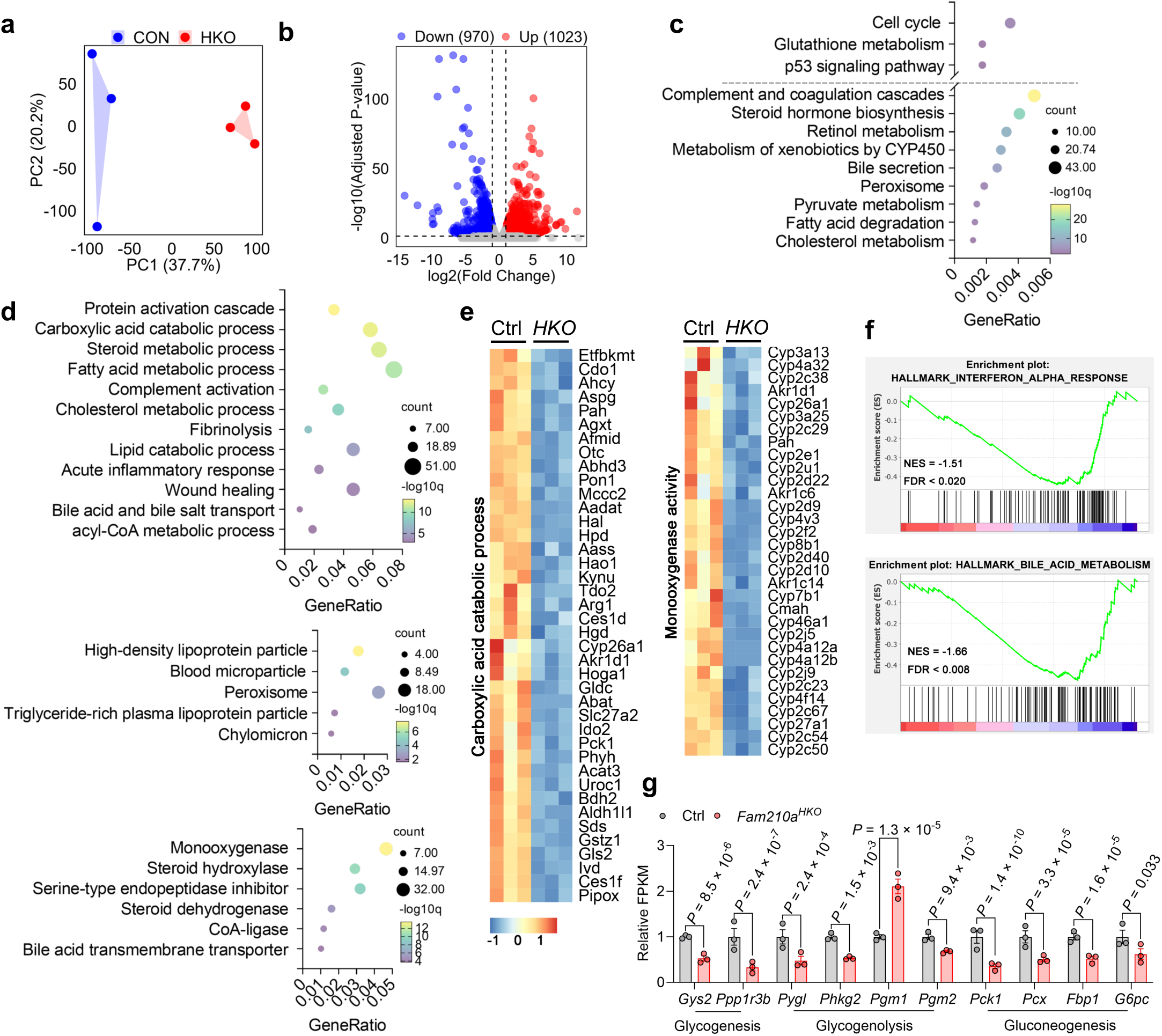
Loss of *Fam210a* impairs hepatic metabolism and function. **a** Principal component analysis (PCA) of liver RNA-seq data from 5-week-old Ctrl and *Fam210a^HKO^* mice. **b** Volcano plot showing differentially expressed genes (DEGs) between Ctrl and *Fam210a^HKO^* livers. **c** KEGG pathway enrichment analysis of significantly downregulated genes. **d** Gene Ontology (GO) enrichment analysis of significantly downregulated genes. **e** Heatmaps of downregulated genes related to carboxylic acid catabolic process and monooxygenase activity. **f** Gene Set Enrichment Analysis (GSEA) identified the enrichments of interferon alpha response and bile acid metabolism downregulated in *Fam210a^HKO^* liver. **g** Expression of genes related to glucose metabolism from the RNA-seq analysis (n = 3 per group; mean ± s.e.m.; t-test).

Moreover, Gene Ontology (GO) analysis of downregulated genes suggests significant enrichments in biological processes including the protein activation cascade, carboxylic acid catabolic process, sterol/cholesterol and fatty acid metabolic processes, acute inflammatory response, and bile acid and bile-salt transport; cellular components including high-density lipoprotein particles and peroxisomes; and molecular functions including monooxygenase activity, steroid-related activities, and bile-acid transporter activity (Fig. 3d, e; Supplementary Fig. 1a). Gene Set Enrichment Analysis (GSEA) further revealed significant negative enrichment of the interferon alpha response and bile acid metabolism gene sets in KO livers relative to controls (NES < 0; FDR < 0.25) (Fig. 3f). These unbiased omics analyses reveal that loss of *Fam210a* disrupts liver function and compromises metabolic integrity.

Motivated by our observation that *Fam210a* KO mice have reduced hepatic glycogen at postnatal 3 and 5 weeks old and lower blood glucose after fast–refeeding, we further looked the changes in genes related to glucose metabolism. RNA-seq data showed that FPKM levels of key genes in glycogenesis (*Gys2* and *Ppp1r3b*), glycogenolysis (*Pygl*, *Phkg2*, and *Pgm2*), and gluconeogenesis (*Pck1*, *Pcx*, *Fbp1*, and *G6pc*) were significantly decreased except the increase of *Pgm1*, indicating broad suppression of hepatic glucose metabolism (Fig. 2g). This profile is consistent with diminished glycogen deposition and, during the late postprandial window, impaired glycogen mobilization and gluconeogenic output, offering a mechanistic explanation for the lower blood glucose in KO mice.

### Impaired mitochondrial respiration and metabolism in juvenile *Fam210a^HKO^* liver

We next sought to determine whether *Fam210a* deletion affects mitochondrial function, sequentially contributing to hepatic dysfunction. To test if FAM210A deficiency alters the mitochondrial proteome of hepatocytes, we first performed proteomic analysis using mitochondria isolated from the livers of 5-week-old control and *Fam210a^HKO^* mice (Fig. 4a). A total of 876 mitochondrial proteins were identified, among which 376 were upregulated and 55 were downregulated in *Fam210a^HKO^* livers (Fig. 4b; Source data 2). GO analysis revealed that the downregulated proteins were significantly associated with responses to oxidative stress together with broad suppression of organic acid, lipid, amide, and cholesterol metabolic pathways, indicating attenuated hepatic redox control and sterol/lipid handling in *Fam210a^HKO^* mice (Fig. 4c, d; Supplementary Fig. 1b). These proteomic changes suggest compromised hepatic redox control and lipid/sterol processing in *Fam210a^HKO^* mice.

**Fig. 4.**
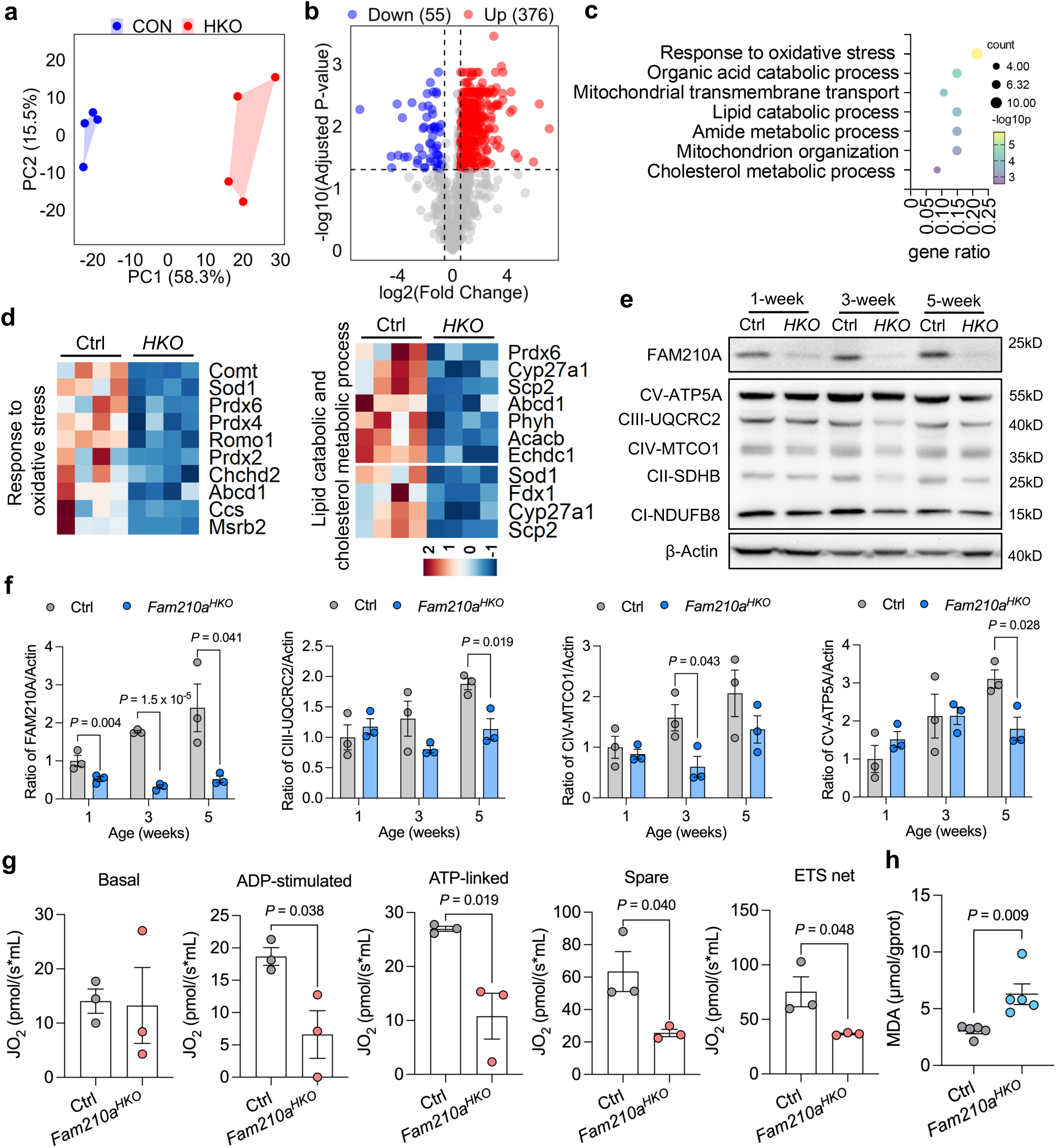
*Fam210a* deletion in hepatocytes impairs liver mitochondrial metabolism and induces lipid peroxidation. **a** PCA of liver mitochondrial proteome profiles from Ctrl and *Fam210a^HKO^* mice. **b** Volcano plot showing differentially expressed proteins between Ctrl and *Fam210a^HKO^* liver mitochondria. **c** GO analysis of significantly downregulated proteins. **d** Heatmaps showing downregulation of proteins related to oxidative stress and lipid metabolism. **e** Western blot analysis of oxidative phosphorylation (OXPHOS) complex proteins in Ctrl and *Fam210a^HKO^* livers at 1, 3, and 5 weeks old. **f** Quantification of relative protein expression in (**e**) (n = 3; mean ± s.e.m.; two-tailed, unpaired Student’s t-test). **g** Mitochondrial respiratory capacity measured with liver mitochondria isolated from 5-week-old mice using the Oroboros O2k (n = 3; mean ± s.e.m.; two-tailed, unpaired Student’s t-test). **h** MDA levels in Ctrl and *Fam210a^HKO^* livers at 3 weeks old (n = 5; mean ± s.e.m.; two-tailed, unpaired Student’s t-test).

To examine the mitochondrial consequences of *Fam210a* loss, we assessed OXPHOS subunits and respiratory capacity. Immunoblotting showed a significant reduction of CIV subunit MT-CO1 at 3 weeks and decreases in CIII subunit UQCRC2 and CV subunit ATP5A at 5 weeks in *Fam210a^HKO^* livers (Fig. 4e, f). Consistently, respiration of purified liver mitochondria was impaired, with ADP-stimulated, ATP-linked, spare, and electron transport chain (ETS) capacities all significantly reduced in KO groups relative to controls (Fig. 4g). These data indicate mitochondrial functional and metabolic deficits upon *Fam210a* deletion. In contrast, α-ketoglutarate dehydrogenase (α-KGDH) activity was unchanged (Supplementary Fig. 1e), suggesting that the primary defect is unlikely to lie in matrix dehydrogenase activity but rather in the abundance and/or assembly of respiratory chain complexes. As a marker of lipid peroxidation, malondialdehyde (MDA) was elevated (Fig. 4h), whereas total glutathione (GSH), glutathione disulfide (GSSG), and the GSH/GSSG ratio were not significantly altered (Supplementary Fig. 1d), indicating maintained global hepatic glutathione homeostasis. Together, *Fam210a^HKO^* livers exhibit impaired mitochondrial respiration and metabolism in juvenile.

### FAM210A regulates mitochondrial ultrastructure and OPA1 processing through interaction with YME1L

To determine whether the observed respiratory defects were associated with altered mitochondrial morphology and content, we performed transmission electron microscopy (TEM) analysis and observed abnormal mitochondrial ultrastructure in *Fam210a^HKO^* livers, characterized by mitochondria with enlarged, balloon-like structures, increased endoplasmic reticulum sheets, and disorganized cristae architecture (Fig. 5a). Moreover, we measured hepatic mtDNA copy number at different developmental stages and found a significant reduction at 3 and 5 weeks in *Fam210a^HKO^* mice (Fig. 5b). These data suggest alterations in mitochondrial biogenesis or maintenance in the liver in the absence of FAM210A.

**Fig. 5.**
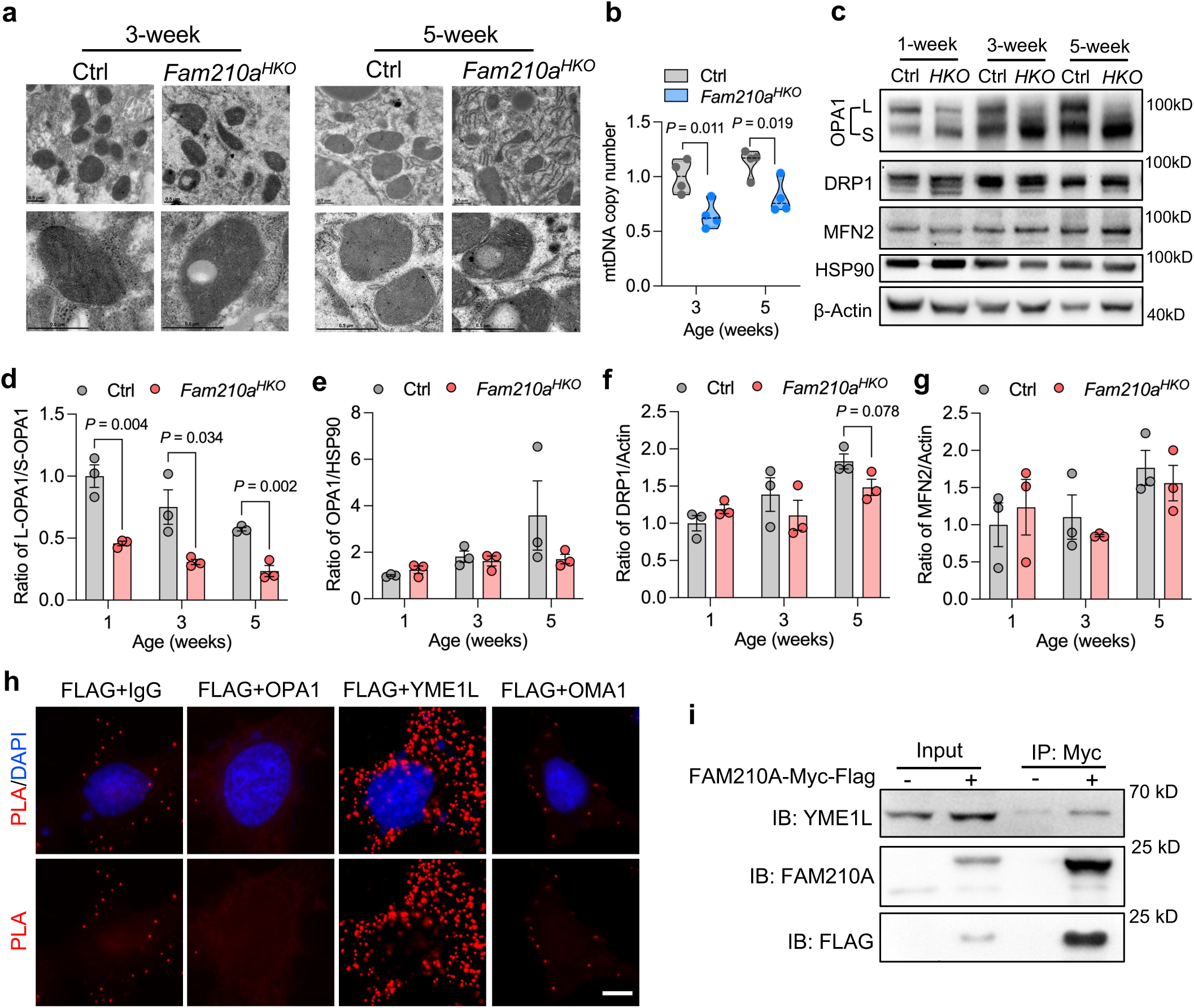
FAM210A deficiency disrupts mitochondrial integrity and promotes YME1L-dependent OPA1 cleavage in hepatocytes. **a** Representative TEM images of liver mitochondria in Ctrl and *Fam210a^HKO^* mice at 3 and 5 weeks old (Scale bar, 0.5 µm). **b** Relative mitochondrial DNA (mtDNA) copy number in the liver of Ctrl and *Fam210a^HKO^* mice at 3, and 5 weeks old (n = 4; mean ± s.e.m.; two-tailed, unpaired student’s *t*-test). **c** Western blot analysis of mitochondrial dynamics–related proteins in Ctrl and *Fam210a^HKO^* livers at 1, 3, and 5 weeks old. **d-g** Quantification of the ratio of L-OPA1/S-OPA1 (**d**), total OPA1 (**e**), DRP1 (**f**), and MFN2 (**g**) protein levels shown in (**c**) (n = 3; mean ± s.e.m.; two-tailed, unpaired student’s *t*-test). **h** Proximity ligation assay (PLA) between FAM210A-FLAG and OPA1, YME1L and OMA1 in *Fam210a-flag* overexpressed HepG2 cells (Scale bar, 5 µm). **i** Co-immunoprecipitation (Co-IP) of FAM210A-Myc and YME1L in *Fam210a-myc* overexpressed HepG2 cells (n = 3 independent experiments).

Given the observed structural abnormalities, we examined the expression of proteins involved in the regulation of mitochondrial dynamics. Strikingly, a remarkable change in OPA1 processing was observed, with a significant decrease in long isoform OPA1 (L-OPA1) and an increase in the short isoform OPA1 (S-OPA1), resulting in the reduced ratio of L-OPA1/S-OPA1 (Fig. 5c, d). In contrast, total OPA1, DRP1, and MFN2 protein levels remained unchanged (Fig. 5e-g). Therefore, FAM210A deficiency dysregulates mitochondrial inner membrane remodeling in hepatocytes by influencing OPA1 processing. We have previously reported that FAM210A interacts with a mitochondrial i-AAA protease YME1L to regulate OPA1 processing in brown adipose tissue^29^. We then sought to test whether this regulatory axis is conserved in the liver and responsible for the mitochondrial remodeling. The proximity ligation assay (PLA) in HepG2 cells showed clear PLA puncta between FAM210A-FLAG and YME1L, instead of the zinc metallopeptidase OMA1 and OPA1 (Fig. 5h), suggesting the potential interaction between FAM210A and YME1L. Further co-immunoprecipitation (Co-IP) assay confirmed the physical interaction between FAM210A and YME1L in HepG2 cells (Fig. 5i). Thus, consistent with the findings in adipocytes, FAM210A regulates mitochondrial cristae organization and OPA1 processing in hepatocytes through interaction with YME1L.

### Hepatic injury caused by *Fam210a* deficiency is resolved in adulthood

Although *Fam210a* deletion impaired mitochondrial structure/function and perturbed hepatic lipid metabolism during juvenile stages, these defects proved transient. By 8 weeks of age, *Fam210^HKO^*mice were indistinguishable from controls in body weight and composition, and liver weight, and liver morphology was comparable to that of control mice (Fig. 6a-d). H&E and BODIPY staining showed no excess lipid deposition in *Fam210^HKO^*livers (Fig. 6e), and PAS staining revealed similar glycogen storage between the livers of *Fam210a^HKO^* and controls (Fig. 6f, g). Moreover, fast–refeeding, GTT, and insulin tolerance test (ITT) demonstrated no differences in blood glucose levels between *Fam210a^HKO^*and controls (Fig. 6h-j), suggesting the indistinct glucose metabolism in adult liver. These findings indicate that the juvenile liver injury caused by *Fam210a* deficiency has resolved by adulthood.

**Fig. 6.**
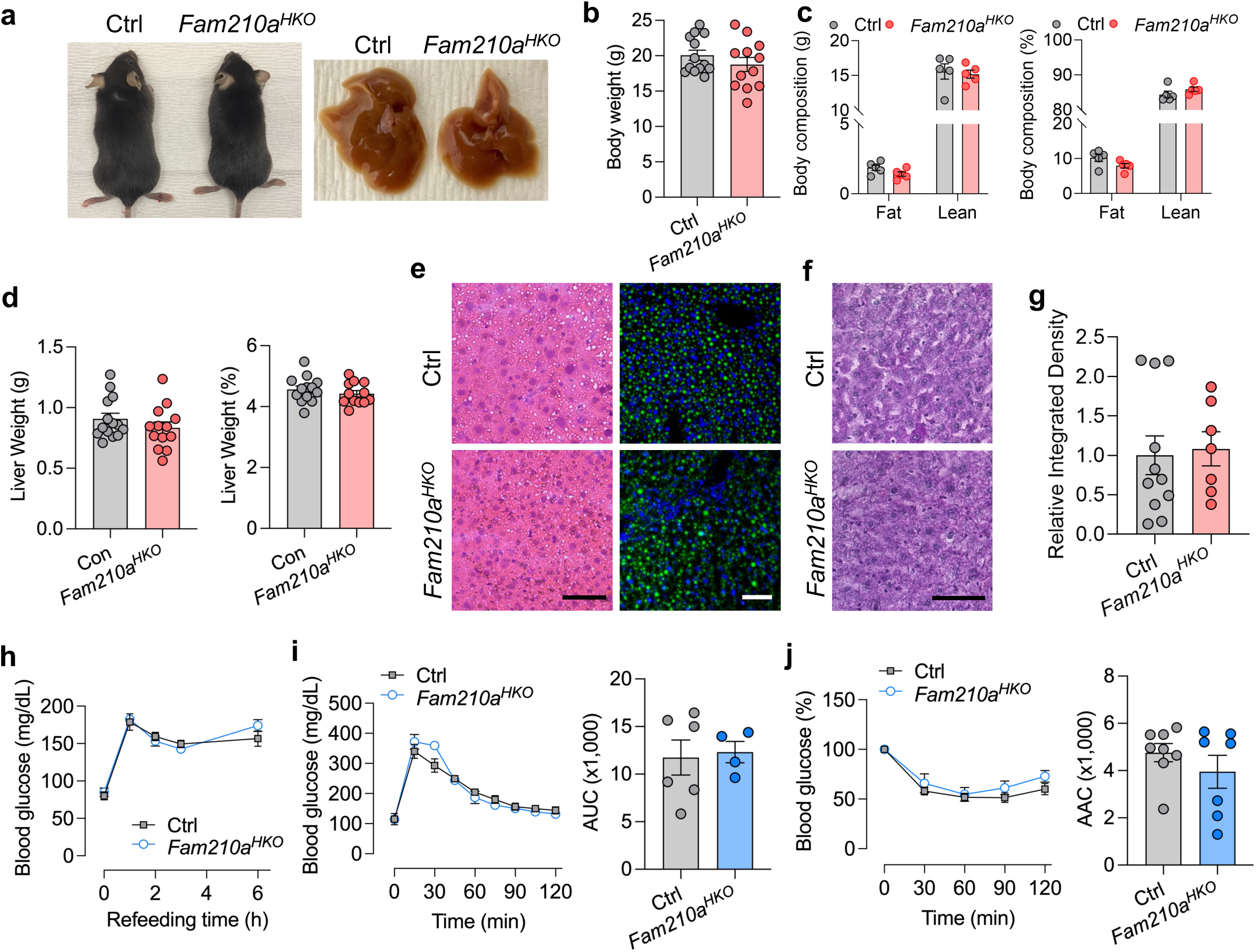
Adult *Fam210a* KO mice are metabolically comparable to control mice. **a** Representative images showing Ctrl and *Fam210a^HKO^* mice at 8 weeks old. **b** Body weight from Ctrl and *Fam210a^HKO^* mice at 8 weeks old (Ctrl, n = 13; *Fam210a^HKO^,* n = 12; mean ± s.e.m.; two-tailed, unpaired student’s *t*-test). **c** Body composition from Ctrl and *Fam210a^HKO^* mice (n = 5; mean ± s.e.m.; two-tailed, unpaired student’s *t*-test). **d** Liver weight and percentage of liver weight from Ctrl and *Fam210a^HKO^* mice (Ctrl, n = 13; *Fam210a^HKO^,* n = 12; mean ± s.e.m.; two-tailed, unpaired student’s *t*-test). **e** Representative H&E and BODIPY staining of liver sections. (Scale bar, 50 µm). **f** Representative PAS staining of liver sections (Scale bar, 50 µm). **g** Quantification of relative integrated density in (**f**) (Ctrl, n = 11; *Fam210a^HKO^,* n = 7; mean ± s.e.m.; two-tailed, unpaired student’s *t*-test). **h** Blood glucose levels in 8-week-old mice after fast-refeeding (n = 7; mean ± s.e.m.; two-tailed, unpaired student’s *t*-test). **i** GTT analysis and area under curve (AUC) of GTT in 5-week-old mice (Ctrl, n = 6; *Fam210a^HKO^,* n = 4; mean ± s.e.m.; two-tailed, unpaired student’s *t*-test). **j** ITT analysis and area above curve (AAC) of ITT in 5-week-old mice (Ctrl, n = 8; *Fam210a^HKO^,* n = 7; mean ± s.e.m.; two-tailed, unpaired student’s *t*-test).

### *Fam210a* deficiency triggers adaptive stress response and promotes hepatic repair

We next aimed to understand why the metabolic disruption and steatosis in *Fam210a^HKO^*mice are resolved in the adult liver. Intriguingly, GO analysis of our RNA-seq data revealed that genes related to chromosome segregation, nuclear division, spindle organization, and positive regulation of cell cycle pathways, were predominantly upregulated enriched in *Fam210a^HKO^* livers (Fig. 7a). The upregulated set spans kinetochore assembly, spindle dynamics, and spindle-assembly checkpoint factors (e.g., *Ndc80*, *Cenp*, *Bub1b*, *Kifs*), indicating enhanced cell-cycle/mitotic pathways (Fig. 7b). RT–qPCR showed that transcripts of the proliferation markers *Cdc20* and *Mki67* were significantly increased in *Fam210a^HKO^* livers at postnatal week 5 (Fig. 7c). Consistently, Ki-67 immunostaining revealed a higher fraction of proliferating (Ki67⁺) cells in the *Fam210a^HKO^*livers than in controls at week 5. The presence of dividing nuclei in the *Fam210a* KO liver sections further indicates active cellular proliferation (Fig. 7e). Beyond cell cycle genes observed in RNA-seq, genes related to ribosome-associated categories were upregulated (Fig. 7f). Notably, GSEA further revealed activation of the unfolded protein response (UPR), and the ISR targets *Atf4*, *Fgf21*, and *Gdf15* were robustly increased in *Fam210a^HKO^* livers, consistent with engagement of the eIF2α–ATF4 axis (Fig. 7g, h). Concordantly, mitochondrial proteome revealed enrichment of translation and organization modules, consistent with UPRmt-like remodeling that supports protein synthesis and structural maintenance (Fig. 7i). GSEA also indicated increased glycolysis and cholesterol homeostasis, reflecting compensatory energy production and membrane remodeling (Supplementary Fig. 2a). Mitochondrial transport and amino acid metabolism pathways were elevated (Fig. 7i; Supplementary Fig. 2b), supporting ATF4-linked substrate supply and anaplerotic fueling of the TCA cycle. Proteins linked to cellular respiration were upregulated, indicating recovery of OXPHOS capacity (Fig. 7i; Supplementary Fig. 2b). In addition, transient TNFα–NF-κB signaling was enriched (Supplementary Fig. 2a), consistent with stress-associated regeneration. Together, these data suggest that mild mitochondrial stress activates ISR/UPRmt pathways to coordinate proteostasis, metabolic rewiring, and hepatic repair.

**Fig. 7.**
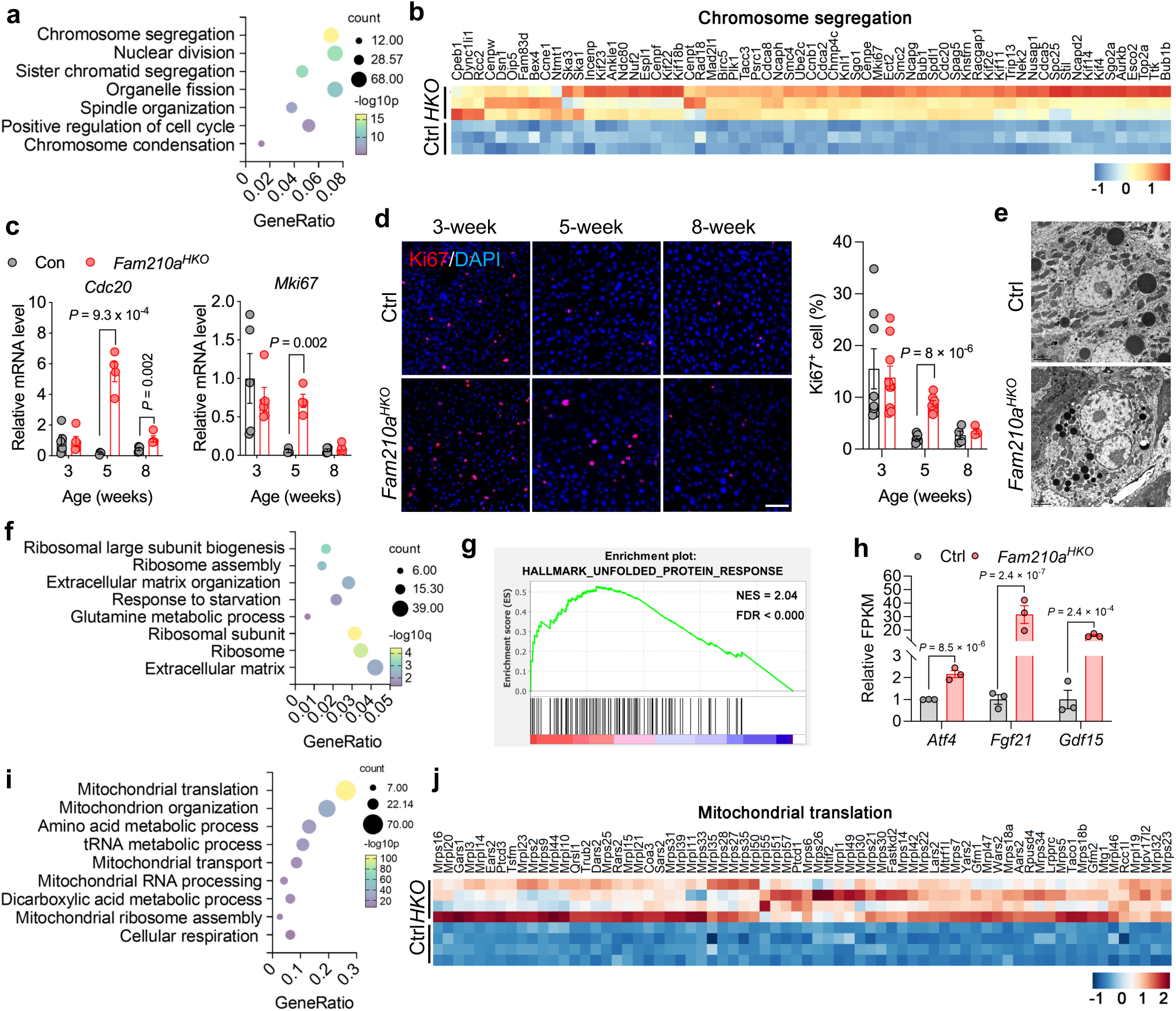
Loss of *Fam210a* induces liver repair. **a** GO analysis of upregulated genes related with cell cycle in the liver from 5-week-old Ctrl and *Fam210a^HKO^* mice. **b** Heatmap showing the expression of proliferation-related genes in Ctrl and *Fam210a* KO livers. **c** Expression levels of representative proliferation-related genes at 3, 5, and 8 weeks of age (n = 5; mean ± s.e.m.; two-tailed, unpaired student’s *t*-test). **d** Immunofluorescence staining of Ki67 and quantification of Ki67^+^ cells in liver sections from control and *Fam210a^HKO^* mice at 3, 5, and 8 weeks old (3-week-old: Ctrl, n = 8; *Fam210a^HKO^*, n = 9; 5-week-old: Ctrl, n = 5; *Fam210a^HKO^*, n = 7; 8-week-old: Ctrl, n = 4; *Fam210a^HKO^*, n = 3; mean ± s.e.m.; two-tailed, unpaired student’s *t*-test, scale bar, 50 µm). **e** TEM images showing a dividing nucleus in the *Fam210a* KO liver at 5 weeks old (Scale bar, 2 µm). **f** GO analysis of upregulated genes in the liver from 5-week-old Ctrl and *Fam210a^HKO^* mice (n = 3). **g** Gene Set Enrichment Analysis (GSEA) identified the enrichment of unfolded protein response upregulated in *Fam210a^HKO^* liver (n = 3). **h** Expression of genes related to adaptive mitochondrial stress response from the RNA-seq analysis (n = 3; mean ± s.e.m.; t-test). **i** GO analysis of upregulated proteins in the liver mitochondria isolated from 5-week-old Ctrl and *Fam210a^HKO^* mice (n = 4). **j** Heatmap showing the expression of mitochondrial translation proteins in Ctrl and *Fam210a* KO livers (n = 4).

### Restoration of mitochondrial and hepatic homeostasis in adult *Fam210a^HKO^* mice

We then asked whether the adaptive responses would restore mitochondrial function in adulthood. Indeed, mtDNA copy number was comparable between adult *Fam210a^HKO^* and control mice (Fig. 8a). Western blot analysis showed that compared to the controls, OPA1 processing had reverted to control-like patterns, and steady-state OXPHOS complex abundance was indistinguishable (Fig. 8b-e). MFN2 remained selectively reduced in adulthood despite normalization of other mitochondrial dynamics markers and respiration (Fig. 8b, c). In line with the mitochondrial structural/functional restoration, genes *Elovl3*, *Slco1a1*, and *Cyp4a12a*, which ranked among the top downregulated transcripts at week 5 and confirmed marked suppression at week 5, showed no differences at week 8 (Fig. 8f). Likewise, complement effector genes (*C8a*, *C8b*, *C9*) that were reduced at week 5 were no longer altered at week 8 (Fig. 8g). Together with equivalent liver and body weights and indistinguishable histology at this time point, these data indicate that the adult livers of *Fam210a*-deficient mice have completed repair and re-established mitochondrial and hepatic homeostasis.

**Fig. 8.**
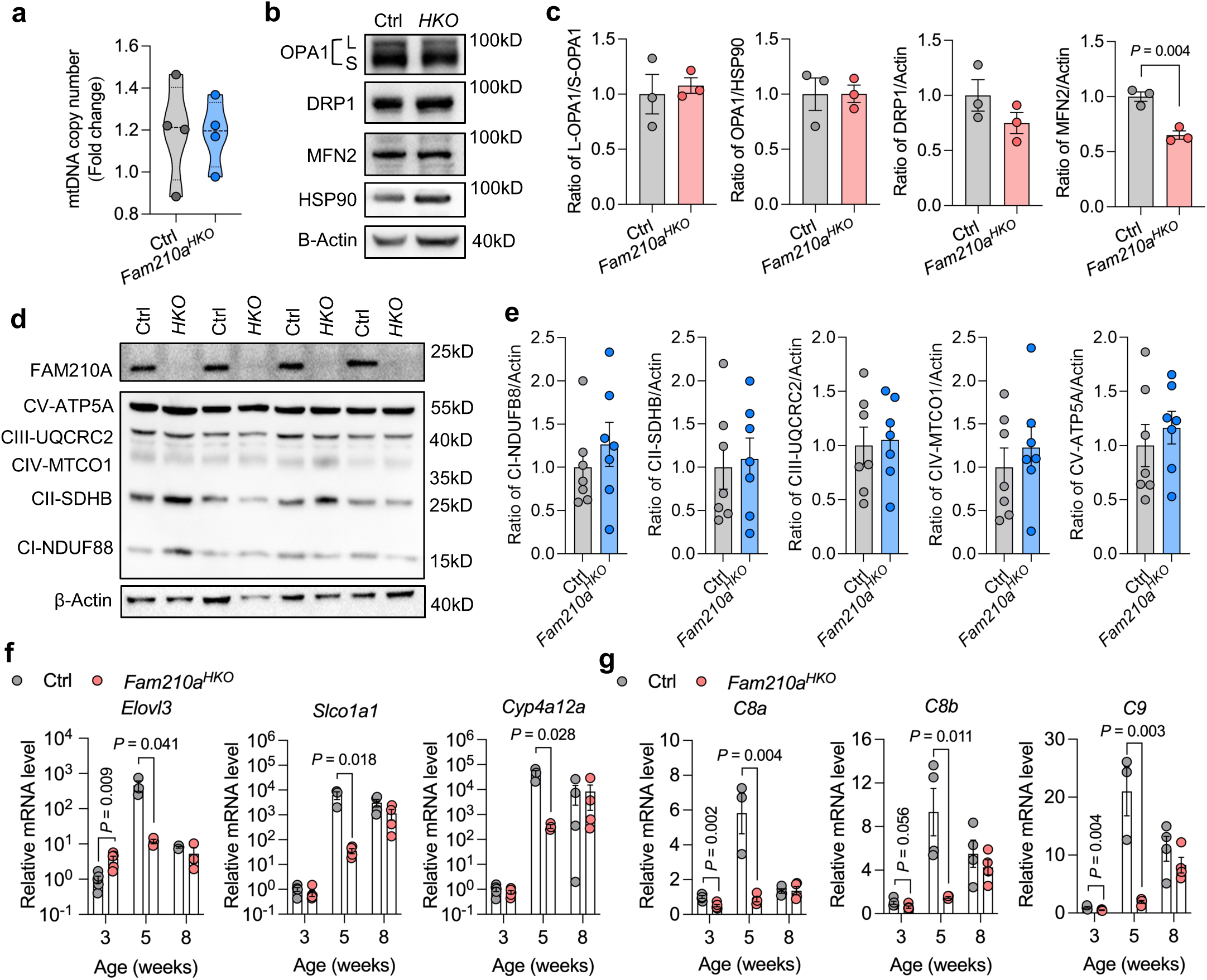
Adult *Fam210a^HKO^* mice exhibit recovery of mitochondrial and hepatic function. **a** Relative mitochondrial DNA (mtDNA) copy number in the liver of Ctrl and *Fam210a^HKO^* mice at 8 weeks old (n = 4; mean ± s.e.m.; two-tailed, unpaired student’s *t*-test). **b** Western blot analysis of mitochondrial dynamics– related proteins in Ctrl and *Fam210a^HKO^* livers. **c** Quantification of relative protein expression in (**b**) (n = 3; mean ± s.e.m.; two-tailed, unpaired student’s *t*-test). **d** Western blot analysis of oxidative phosphorylation (OXPHOS) complex proteins in the liver from Ctrl and *Fam210a^HKO^* mice at 8 weeks old. **e** Quantification of relative protein expression in (**d**) (n = 7; mean ± s.e.m.; two-tailed, unpaired student’s *t*-test). **f, g** Expression levels of representative hepatic metabolic (**f**) and complement-related (**g**) genes in livers at 3, 5, and 8 weeks old (Ctrl, n = 3; *Fam210a^HKO^*, n = 4; mean ± s.e.m.; two-tailed, unpaired student’s *t*-test).

## Discussion

Postnatal liver development involves the transition from immature hepatoblast-like cells to fully mature hepatocytes, a process that requires both rapid proliferation and extensive metabolic reprogramming^32,37,38^. Single-cell transcriptomic studies show that hepatocytes remain immature through the first three weeks, gradually acquiring mature metabolic features by 7–8 weeks^39^. This maturation is critically dependent on intact mitochondrial remodeling, which coordinates energy production and biosynthetic functions to support hepatic morphogenesis and competence^39^. Yet, the precise mitochondrial mechanisms guiding this process remain incompletely defined. Our findings identify FAM210A as a key regulator of this developmental remodeling, linking mitochondrial structure to hepatic metabolic capacity. Moreover, the discovery of a transient juvenile vulnerability and subsequent compensatory repair highlights a stress-adaptive program with potential therapeutic relevance for liver injury and diseases.

It has been well established that mitochondrial dynamic regulators such as OPA1, MFN1/2, and DRP1 orchestrate fusion, fission, and organelle tethering to maintain mitochondrial structure and bioenergetics^40–43^. Our findings add FAM210A to this foundation, highlighting its indispensable role in sustaining mitochondrial integrity across diverse tissues. Previous work has demonstrated that FAM210A is required in brown adipose tissue to support cold-induced mitochondrial remodeling through OPA1 processing, in cardiomyocytes to promote mitochondrial translation and contractile function, and in skeletal muscle to couple cytosolic protein synthesis with mitochondrial metabolism^25,28,29^. Extending these findings, we show that in the liver, FAM210A preserves mitochondrial cristae integrity and oxidative phosphorylation capacity by regulating YME1L-dependent OPA1 processing. Hepatocyte-specific deletion of *Fam210a* disrupted ultrastructure, reduced mtDNA abundance, and impaired respiratory function, resulting in juvenile hepatic steatosis, hypoglycemia, and systemic metabolic imbalance. Notably, unlike its role in brown fat, heart, or skeletal muscle where FAM210A deficiency causes progressive dysfunction, hepatic loss of FAM210A triggered a transient hepatopathy that resolved in adulthood through compensatory hepatocyte proliferation, mitochondrial biogenesis, and integrated stress response activation. These findings underscore both the shared requirement of FAM210A for mitochondrial integrity across tissues and its unique role in the liver as a developmental gatekeeper of metabolic maturation and a trigger of adaptive remodeling programs.

Our findings show that *Fam210a* deficiency in the liver causes hypoglycemia after fasting– refeeding, reduced glycogen storage, and suppression of genes for glycogenesis, glycogenolysis and gluconeogenesis, indicating defective hepatic glucose storage and mobilization. Although the precise mechanisms remain to be defined, we predict that FAM210A regulates glucose metabolism indirectly through its role in maintaining mitochondrial structure and respiration. Loss of FAM210A may limit ATP and NADH supply for gluconeogenesis, disrupt TCA cycle flux and acetyl-CoA availability for glycogen synthesis, and impair metabolite signaling such as acetylation and redox balance that controls transcription of glucose metabolic genes^28^. Notably, knockout of other mitochondrial dynamics genes such as *Opa1* or *Drp1* alters glucose tolerance and insulin sensitivity^44–46^, whereas *Fam210a* deletion did not significantly affect GTT or ITT under basal conditions. It will therefore be important to test whether glucose tolerance, systemic insulin sensitivity, and energy expenditure become altered in FAM210A-deficient mice under conditions of diet-induced obesity or experimental liver disease, which may unmask metabolic vulnerabilities.

Our data suggest that FAM210A regulates OPA1 processing through its interaction with YME1L. However, the precise mechanism by which FAM210A maintains mitochondrial integrity in the liver remains incompletely defined. This observation raises an intriguing possibility that modulation of YME1L activity might rescue the mitochondrial and metabolic defects observed in juvenile *Fam210a*-deficient livers. Nevertheless, *in vivo* rescue experiments directly testing this hypothesis remain to be performed. It has been reported that OPA1 deficiency affects mitochondria–ER and peroxisome networks^45^. Intriguingly, we observed enrichment of peroxisome pathways and increase of ER in *Fam210a* KO liver. Whether FAM210A is involved in regulating the mitochondria–ER and peroxisome network warrants further investigation. Previous studies in other tissues have shown that FAM210A can regulate mitochondrial protein translation via interaction with EF-Tu^25^, but we did not observe reductions in mitoproteins or EF-Tu abundance in the liver. Instead, mitoribosomal proteins were upregulated during the recovery phase, indicating enhanced mitochondrial translation as part of the repair process. An additional mechanism may involve regulation of TCA cycle flux and acetyl-CoA metabolism, which influences protein acetylation and has been implicated in skeletal muscle FAM210A deficiency^28^. Whether similar pathways contribute to the hepatic phenotype is worth testing. Furthermore, the stress-adaptive response in *Fam210a* KO livers may involve FAM210A-dependent regulation of YME1L toward OMA1 degradation. Since OMA1 is a key activator of the ISR through DELE1 signaling^47–49^, the absence of FAM210A could destabilize this axis, thereby engaging integrated stress pathways to support recovery. Although not directly tested in this study, these potential mechanisms highlight FAM210A as a multifaceted regulator of mitochondrial proteostasis and signaling, warranting deeper mechanistic interrogation in future work.

Recent studies underscore the critical role of mitochondrial dynamics and lipid remodeling in a variety of chronic liver diseases including alcohol-associated liver disease (ALD), metabolic dysfunction–associated steatohepatitis (MASH)^10,11,14,15,50^. Hepatocyte-specific OPA1 deficiency induces a mitohormetic stress response that protects against drug-induced liver injury and reduces lipid absorption by disrupting mitochondria–peroxisome–ER tethering, thereby mitigating steatosis under high-fat diet conditions^44,45^. By contrast, MFN2 loss abolishes ER–mitochondria phosphatidylserine transfer and precipitates a NASH-like phenotype and liver cancer^51^, while DRP1 dysregulation has context-dependent consequences ranging from protection against NAFLD to exacerbation of alcoholic hepatitis through ER stress and hepatocyte death^52–56^. Similarly, aberrant cardiolipin remodeling can either prevent steatosis or drive spontaneous MASH with fibrosis, underscoring the fine balance between protective adaptation and maladaptive injury^57,58^. In this context, our findings demonstrate that the transient mitochondrial dysfunction and steatosis caused by FAM210A deficiency can be compensated through mitohormetic stress responses and regenerative programs. This raises the important possibility that *Fam210a* deletion–induced adaptive remodeling may enhance hepatic resilience and protect against drug-induced liver injury or diet-induced steatohepatitis. Defining the mechanisms by which FAM210A loss activates these adaptive pathways will be critical, as they may represent exploitable therapeutic strategies to mitigate steatosis, fibrosis, and hepatocellular injury in chronic liver disease.

## Materials and methods

### Ethical statement

This study complied with all relevant ethical regulations. All animal procedures were conducted in accordance with National Institutes of Health and Institutional guidelines and were approved by the Institutional Animal Care and Use Committee (IACUC) at the University of Florida.

### Animals

All mice used in this study were on a C57BL/6J genetic background. The *Fam210a^flox/flox^* mice were generated using the CRISPR-Cas9 system as previously described^29^. *Albumin-Cre* (*Alb^Cre^*, JAX stock #003574) mice were obtained from The Jackson Laboratory (The Jackson Laboratory, Bar Harbor, ME). Genomic DNA was extracted from mouse tail tissue for genotyping by PCR, following the protocols and primer sequences provided by the supplier. The genotypes of experimental knockout and corresponding control mice are as follows: *Fam210a^HKO^* (*Alb^Cre^*; *Fam210a^flox/flox^*) and control (*Fam210a^flox/flox^*). Mice were housed and maintained in the animal facility with free access to water and standard rodent chow food (2018 Teklad Global 18% Protein Rodent Diets), and were housed under 12-h light/dark cycle, at 22 °C, and 45% humidity on average. For all animal-based experiments, at least three pairs of gender-matched littermates at the age of 2–5 months were used. Mice were euthanized using 100% medical grade carbon dioxide followed with cervical dislocation.

### *In vitro* cell culture

HepG2 cell line (a gift from Dr. Mingyi Xie, University of Florida) was cultured in MEM (Corning, cat# MT10009CV) supplemented with 10% Fetal Bovine Serum (FBS, Corning, cat# 35-011-CV) and 1% penicillin/streptomycin (P/S, Gibco, cat# 15-140-122) at 37 °C with 5% CO_2_, followed by replacing with fresh culture medium every two days.

### Blood glucose measurement

For the glucose tolerance test (GTT), mice were fasted overnight and then injected intraperitoneally with D-glucose (2 g/kg body weight). Blood glucose levels were measured at indicated time points from the tail vein using a glucometer (Accu-Check Active, Roche). For the insulin tolerance test (ITT), mice were fasted for 4 hours prior to intraperitoneal injection of insulin (ProZinc) (0.75 U/kg body weight), and tail blood glucose was similarly monitored over time. For the fasting-refeeding experiment, mice were fasted overnight, and blood glucose levels were measured at 0, 1, 2, 4, and 6 hours after refeeding.

### Body composition assessment

Body composition was measured using an EchoMRI whole body composition analyzer (EchoMedical Systems). The instrument was calibrated on each day of use with manufacturer-supplied canola-oil and phantom standards. Mice were weighed immediately before scanning. Then each mouse was placed in a ventilated, size-appropriate plastic restraint tube and scanned. Outputs included fat mass and lean mass.

### Mitochondrial isolation

Mitochondria were isolated from the liver tissue of mice at 5 weeks old following the protocol previous reported with modifications^59^. Briefly, 100 mg liver tissue was washed with cold PBS, mince into pieces, and homogenized with a Cole-Parmer PTFE tissue grinder in 2 mL isolation buffer (225 nM Mannitol, 75 nM Sucrose, and 0.2 nM EDTA in distilled water, pH 7.4) for 8-10 strokes. The homogenate was centrifuged at 1000 × g for 10 min at 4 °C, and the top layer containing fat was discarded. The supernatant was transferred into a new tube and centrifuged at 6200 × g for 10 min at 4 °C. The supernatant was discarded, and the pellet was resuspended in 1 mL isolation buffer, and centrifuged at 6200 × g for 10 min at 4 °C. The sediment mitochondria were resuspended in 100 μL isolation buffer, and the protein concentration was determined using Pierce BCA Protein Assay Reagent (Pierce Biotechnology, cat# 23225).

### Oxygen consumption rate (OCR) measurement

Mitochondria were isolated from livers of 5-week-old mice as described above, and then resuspended in respiration medium (Mir05: 0.5 mM EGTA, 3 mM MgCl₂, 60 mM K-lactobionate, 20 mM taurine, 10 mM KH₂PO₄, 20 mM HEPES, 110 mM sucrose, 1 g·L⁻¹ fatty-acid–free BSA; pH 7.1). High-resolution respirometry was performed using the Oxygraph-2k (Oroboros Instruments) to assess ADP-stimulated respiration, ATP-linked respiration, spare respiratory capacity, and ETS capacity. A substrate-inhibitor titration protocol was performed according to DL-Protocol: Suit-006_O2_mt_D047 in 0.5mL chambers. Briefly, 20 µL of mitochondrial suspension (0.04 mg/mL) was added to a 0.5 mL O2k chamber (MiR05, 37 °C, 750 rpm). Substrates and effectors were introduced at the following final concentrations in sequence: 5 mM pyruvate and 2 mM malate to record baseline LEAK; 2.5 mM ADP/Mg²⁺ to obtain OXPHOS capacity; 10 µM cytochrome c to verify outer-membrane integrity; 5 nM oligomycin to define LEAK with ATP-synthase inhibited; 0.5 µM CCCP titrated in three steps to reach peak ETS capacity; and 2.5 µM antimycin A to determine residual oxygen consumption.

### Alpha ketoglutarate dehydrogenase complex (α-KGDH) activity assay

The enzymatic activity of α-ketoglutarate dehydrogenase (α-KGDH) was measured fluorometrically by monitoring NADH autofluorescence (excitation/emission = 340/450 nm) in black 96-well plates using a BioTek Synergy 2 Multimode Microplate Reader. The reaction was performed in KGDH Assay Buffer, consisting of 105 mM K-MES, 30 mM KCl, 1 mM EGTA, 10 mM K₂HPO₄, 5 mM MgCl₂·6H₂O, and 2.5 mg/mL BSA (pH 7.2). The buffer was supplemented with 0.03 mg/mL alamethicin, 5 μM rotenone, and 2 mM NAD⁺ to support mitochondrial membrane permeability and electron transport inhibition. Additionally, 0.1 mM coenzyme A and 0.3 mM thiamine pyrophosphate were included as essential cofactors for α-KGDH activity. After adding the assay buffer to each well, isolated mitochondria were added, and the reaction was initiated by the addition of 10 mM α-ketoglutarate. The rate of NADH production was calculated from the slope of the linear portion of the fluorescence curve. Fluorescence intensity was converted to picomoles of NADH using a standard curve generated with known concentrations of NADH.

### Vector construction and cell transfection

The pCMV6-Fam210a-Myc-DDK plasmid was obtained from our previous study^29^. The plasmid was transfected into HepG2 cells using PEI MAX (Kyfora Bio, cat# 24765). Cells were subjected to proximity ligation assay or harvested for co-immunoprecipitation 48 hours post-transfection.

### Transmission electron microscopy (TEM)

For TEM imaging, samples were dissected and cut in 1 mm^3^ pieces that were immediately immersed into Trumps fixative and kept at 4 °C for 48 hours. The fixed samples were placed in fresh fixative followed by several buffer washes with 0.1 M sodium cacodylate, pH 7.24, post-fixed with buffered 2% OsO_4_, water washed and dehydrated in a graded ethanol series (25%-100% in 5% increments) followed by 100% acetone. Dehydrated samples were infiltrated with Embed/Araldite epoxy resin and Z6040 embedding primer (EMS, Hatfield, PA) and cured for 72 hours at 65–75 °C. Ultra-thin sections were collected on carbon coated Formvar 100 mesh copper grids (EMS, Hatfield, PA) and post-stained with 2% aqueous uranyl acetate and Reynold’s lead citrate. Sections were examined with a Talos L120C G2 TEM (Thermo Fisher Scientific) and digital images were acquired with a Ceta 16M pixel CMOS camera and Velox User Interface (Thermo Fisher Scientific) in the ICBR Electron Microscopy Core Facility at the University of Florida.

### Immunofluorescence staining

Immunofluorescence was performed on liver tissue sections. Briefly, liver tissues embedded in Tissue-Tek O.C.T Compound were sectioned at 10 μm thickness using a Microm HM525 cryostat. The samples were fixed in 4% PFA for 20 min, and permeabilized and blocked in blocking buffer (PBS containing 5% goat serum, 2% bovine serum albumin, 0.2% Triton X-100, and 0.1% sodium azide) for 1 h at room temperature (RT). Samples were incubated overnight at 4 °C with primary antibodies diluted in blocking buffer. Following PBS washes, the samples were incubated with secondary antibodies, Bodipy 493/503 (Invitrogen, cat# D3922), and DAPI (Invitrogen, cat# D1306). Antibodies and dilutions were used as follows: Ki67 (AbCam, ab15580, 1:1000) and Alexa 647 goat anti-rabbit IgG (Invitrogen, A-21244, 1:1000).

### H&E and Periodic acid-Schiff (PAS) staining

For H&E staining, cryosections from control and *Fam210a^HKO^* mice were fixed in 95% EtOH for 2 min. The sections were stained with hematoxylin for 15 min, then rinsed in running tap water for 5 min and stained with eosin for 1 min. Slides were dehydrated through 70%, 90%, 95%, and 100% ethanol (10 s each), cleared in xylene for 2 min, and mounted with glass coverslips using Permount. For PAS staining, cryosections were stained with periodic acid for 5 min and rinsed in running tap water for 3 min, then strained with Schiff reagent for 5 min and rinsed in running tap water for 3 min. Slides were then dehydrated through 70%, 70%, 96%, 96%, 100%, and 100% ethanol (1 min each), cleared in xylene (2 changes, 5 min each), and then covered using Permount for further imaging.

### DNA extraction and mitochondria DNA copy number analysis

Liver tissue was digested in 500 μL DNA extraction buffer (10 mM Tris-HCL (pH 8.0), 1% SDS, 50 mM EDTA, 20 mM NaCL, 50 μg Proteinase K) overnight at 55 °C. After digestion, 500 μL phenol was added to the sample, followed by centrifugation at 12,000 for × g for 20 min at 4 °C. The supernatant containing DNA was transferred to a new tube, an equal volume of phenol: chloroform: Isoamyl solution (25:24:1, v/v) was added, followed by centrifugation at 12,000 × g for 15 min at 4 °C. The same volume of chloroform: isoamyl alcohol (24:1, v/v) was added to the aqueous phase, followed by centrifugation at 10,000 × g for 15 min. The upper phase was carefully transferred to a new tube, and genomic DNA was precipitated by the addition of 1/10 volume of 3 M sodium acetate and 2x volumes of precooled 100% ethanol. The mixture was incubated at −20 °C for 20 min, followed by centrifugation at 10,000 × g for 10 min. After discarding the supernatant, the DNA pellet was resuspended in 30 μL of nuclease-free water. DNA concentration and quality were assessed using NanoDrop 2000c and subjected to subsequent real-time PCR analysis. To quantify the number of mitochondria DNA (mtDNA), we used the following *NADH dehydrogenase subunit 2* (*Nd2*) primers: Forward: CCTATCACCCTTGCCATCAT; Reverse: GAGGCTGTTGCTTGTGTGAC. To quantify nuclear DNA, we used the *Platelet and Endothelial Cell Adhesion Molecule 1* (*Pecam1*) primers: Forward: ATGGAAAGCCTGCCATCATG; Reverse: TCCTTGTTGTTCAGCATCAC.

### Total RNA extraction and real-time PCR

Total RNA was isolated from tissue samples using TRIzol reagent (Sigma-Aldrich, cat# T9424) following the manufacturer’s protocol. To eliminate residual genomic DNA, the RNA was treated with RNase-free DNase I. RNA purity and concentration were assessed using a NanoDrop 2000c spectrophotometer (Thermo Fisher). For cDNA synthesis, 3 μg of total RNA was reverse transcribed with M-MLV reverse transcriptase (Invitrogen, cat# 28025021) using random primers. Quantitative real-time PCR was performed on a CFX Opus 96 Real-Time PCR System (Bio-Rad) using SsoFast EvaGreen Supermix with Low ROX (Bio-Rad, cat# 1725211) and gene-specific primers (listed in Supplementary Table 1). Relative gene expression levels were calculated using the 2^−ΔΔCt^ method^60^, with β-actin serving as the internal reference gene.

### Bulk RNA-seq

RNA sequencing was performed by Novogene with Illumina NovaSeq X Plus platform. The raw mRNA sequencing data were initially assessed for quality using FastQC. Low-quality reads and adapter sequences were trimmed using Trimmomatic. The cleaned reads were then aligned to the reference genome using STAR. Gene expression levels were quantified using featureCounts, and expression data were normalized using TPM (Transcripts Per Million). Differential gene expression analysis was performed using the DESeq2 package in R. Statistical significance was determined using an adjusted p-value threshold of <0.05, with p-values corrected for multiple testing using the Benjamini-Hochberg procedure to control the False Discovery Rate (FDR), and an absolute fold change (|FC|) of ≥2 was considered biologically significant. Functional annotation and pathway enrichment analyses were conducted using DAVID and Gene Set Enrichment Analysis (GSEA) to identify significantly enriched Gene Ontology (GO) terms and biological pathways. Data visualization included generating volcano plots, heatmaps, and Principal Component Analysis (PCA) plots using ggplot2 and pheatmap to illustrate differential expression results and sample clustering.

### Protein extraction and immunoblotting analysis

Total protein was isolated from liver tissues using RIPA buffer (25 mM Tris-HCL (pH 8.0), 150 mM NaCl, 1mM EDTA, 0.5% NP-40, 0.5% sodium deoxycholate, and 0.1% SDS). Protein concentrations were measured using the Pierce BCA Protein Assay Kit. Equal amounts of protein were separated by SDS-PAGE and transferred onto polyvinylidene fluoride (PVDF) membranes (Millipore Corporation). Membranes were blocked with 5% non-fat milk in TBST for 1 hour at RT, followed by overnight incubation at 4 °C with primary antibodies diluted in 5% milk. After washing, membranes were incubated with appropriate HRP-conjugated secondary antibodies for 1 hour at RT. Following antibodies were used: C18orf19 (FAM210A, Invitrogen, PA5-53146; 1:1000), β-Actin (Santa Cruz, sc-47778; 1:1000), OxPhos Rodent WB Antibody Cocktail (Invitrogen, 45-809-9; 1:2000), OPA1 (BD Biosciences, 612606; 1:1000), DRP1 (Proteintech, 12957-1-AP; 1:1000), Mitofusin 2 (MFN2, Cell Signaling, 9482; 1:1000), HSP90 (Proteintech, 13171-1-AP; 1:2000), YME1L (Proteintech, 11510-1-AP; 1:1000), FLAG (Proteintech, 20543-1-AP; 1:20000), HRP AffiniPure goat anti-mouse IgG (Jackson ImmunoResearch, 115-035-003; 1:10000), HRP AffiniPure goat anti-rabbit IgG (Jackson ImmunoResearch, 111-035-003; 1:10000). Immunodetection was performed using Pierce™ ECL Western Blotting Substrate (Fisher Scientific, cat# 32106) and visualized using the iBright 750 Imaging System (Thermo Fisher Scientific). The results shown are representative of at least three independent experiments.

### Sample preparation for mitochondria proteomics

Proteins were extracted from purified mitochondria using the EasyPep™ MS Sample Prep Kit (Thermo Fisher Scientific) and quantified with a Qubit fluorometer. For each sample, an appropriate volume was taken to yield 20 µg total protein. Samples were digested with the sequencing-grade Trypsin/Lys-C Rapid Digestion Kit (Promega, Madison, WI) following the manufacturer’s instructions. Briefly, 3× the sample volume of Rapid Digestion Buffer was added. Samples were reduced with 1 µL DTT (0.1 M in 100 mM ammonium bicarbonate) at 56 °C for 30 min, followed by alkylation with 0.54 µL iodoacetamide (55 mM in 100 mM ammonium bicarbonate) at room temperature in the dark for 30 min. Trypsin/Lys-C was freshly prepared at 1 µg/µL in Rapid Digestion Buffer; 1 µL enzyme was added to each sample, and digestion proceeded at 70 °C for 1 h. Reactions were quenched by adding trifluoroacetic acid to 0.5% (v/v). Peptides were subjected to LC–MS/MS immediately to ensure high-quality tryptic peptides with minimal non-specific cleavage.

### Liquid chromatography (LC)-mass spectrometry MS) analysis

Nano-LC/MS/MS was performed on a Q Exactive HF Orbitrap mass spectrometer (Thermo Scientific) equipped with an EASY-Spray nanospray source (positive-ion mode) and an UltiMate™ 3000 RSLCnano system. Mobile phase A was water with 0.1% formic acid; mobile phase B was acetonitrile with 0.1% formic acid. For loading pump, mobile phase A was water with 0.1% trifluoroacetic acid. 5 µL of each sample was loaded onto a Thermo Scientific µPAC™ C18 trap (pillar diameter 5 µm, length 10 mm, inter-pillar distance 2.5 µm) at 10 µL/min and washed for 3 min at 1% B. Peptides were then eluted to a µPAC™ C18 analytical column (110 cm, 5 µm pillars, 2.5 µm inter-pillar distance) held at 40 °C. The flow was 750 nL/min from 0–15 min and 300 nL/min thereafter. The gradient was 1%→20% B over 100 min, then 20%→45% B over 23 min, followed by a 95% B wash (5 min) and re-equilibration at 1% B, for a total run time of 150 min. Source settings: spray voltage 1.5 kV, capillary temperature 200 °C. Data-dependent acquisition was used (Top15): full MS m/z 375–1575 at 60,000 resolution, AGC 3×10^^^, max IT 50 ms; HCD MS/MS at 15,000 resolution, AGC 2×10^5^, max IT 55 ms, NCE 28, 4 m/z isolation window; z=1 precursors excluded; dynamic exclusion 15 s with isotope exclusion; m/z 445.12003 was used as lock mass. A HeLa digest standard was run to verify performance; if protein IDs fell below 3,200, the instrument was cleaned and columns were replaced.

### Data analysis of proteomics

Raw files were processed in Proteome Discoverer 3.2.0.450 (Thermo Fisher Scientific) using the Chimerys node for peptide/protein identification. Spectra were searched against Mus musculus (sp_incl_isoforms; TaxID 10090 and subtaxonomies, v2023-06-28) plus a universal contaminant FASTA (Frankenfield et al., 2022), with trypsin specificity, carbamidomethyl-Cys as a fixed modification, precursor tolerance 10 ppm, and fragment tolerance 0.020 Da. Label-free quantification used precursor-ion intensities in Proteome Discoverer: samples were organized with a non-nested study factor for the control and *HKO* group; all peptides were used for normalization; protein abundances were computed as summed peptide-group abundances. Statistics used pairwise ratio-based Fisher’s exact tests with low-intensity resampling imputation, and Benjamini–Hochberg adjustment to derive adjusted p-values. Functional enrichment analysis was performed using DAVID and Metascape.

### Proximity ligation assay (PLA)

Proximity ligation assay (PLA) was performed using the Duolink® In Situ Red Starter Kit (Sigma-Aldrich, cat# DUO92101) as described previously^29^. After immunofluorescence-based primary antibody incubation, PLA reactions were carried out in a 20 μl volume per slide per step. Secondary PLA probes, anti-mouse MINUS and anti-rabbit PLUS, were diluted 1:5 in the provided antibody diluent and incubated overnight at 4 °C. Slides were then washed once with wash buffer A for 10 min. For the ligation step, ligation buffer was prepared by diluting ligation stock 1:5 in distilled water and supplementing with ligase at 1:40. Slides were incubated at 37 °C for 1 h, followed by a 5 min wash in buffer A. The amplification mix was prepared by diluting amplification stock 1:5 and polymerase 1:80 in distilled water, and applied to slides for 2 h at 37 °C. After amplification, slides were washed sequentially with 1× wash buffer B (10 min), 0.01× wash buffer B (1 min), and 1× PBS (1 min). Nuclei were counterstained with DAPI, and images were acquired using a Leica THUNDER 3D Imager fluorescence microscope. The following primary antibodies were used: normal rabbit IgG (Cell Signaling, 2729; 1:300), anti-FLAG (Sigma, F1804; 1:300), anti-YME1L (Proteintech, 11510-1-AP; 1:150), anti-OMA1 (Proteintech, 17116-1-AP, 1:200), anti-FLAG (Proteintech, 20543-1-AP, 1:200), and anti-OPA1 (BD BioScience, 612606-18, 1:75).

### Co-immunoprecipitation

Total protein was extracted from *Fam210a-myc-Flag* overexpressed 293T cells using lysis buffer (20 mM Tris-HCL (pH 8.0), 137 mM NaCl, 0.25% NP-40, and 2 mM EDTA). The lysate containing 1 mg total protein was incubated with Pierce Anti-c-Myc Magnetic Beads (Fisher Scientific, cat#88842) at 4 °C overnight. The samples were washed three times and subjected to immunoblotting analysis.

### Hepatic triglyceride measurement

Liver triglyceride (TAG) content was quantified using the Triglyceride Colorimetric Assay Kit (Cayman Chemical, cat# 10010303) following the manufacturer’s instructions. Briefly, ∼10– 20 mg of frozen liver tissue was homogenized in 400 µl of chloroform:isopropanol:igepal (7:11:0.1) using a refrigerated bead-based homogenizer (Benchmark Scientific), followed by centrifugation at 15,000 × g for 10 min at 4 °C. The supernatant was collected, dried under nitrogen, and resuspended in 400 µl of NP-40 Substitute Assay Reagent (1X). To improve lipid solubility, samples were heated to 100 °C for 1 min, cooled, and vortexed, and this step was repeated once. Homogenates were diluted as necessary in NP-40 Substitute Assay Reagent (1X). Ten microliters of each sample were loaded in duplicate onto a 96-well assay plate along with TAG standards. Reactions were initiated by addition of 150 µl of the Triglyceride Enzyme Mixture, incubated at room temperature for 60 min, and absorbance was measured at 540 nm using a microplate reader. TAG concentrations were calculated from the standard curve and normalized to tissue weight.

### Lipid peroxidation assay

Lipid peroxidation in liver tissue was assessed by measuring malondialdehyde (MDA) using the TBARS (TCA Method) Assay Kit (Cayman Chemical, cat# 700870). Approximately 25 mg of frozen liver tissue was homogenized in RIPA buffer containing protease inhibitors, followed by centrifugation at 1,600 × g for 10 min at 4 °C. The supernatant was incubated with trichloroacetic acid (10%) and thiobarbituric acid (TBA) color reagent under acidic conditions at 95 °C for 1 h to generate MDA–TBA adducts. After cooling and centrifugation, 200 µl of the supernatant was transferred to a 96-well plate, and absorbance was measured at 535 nm using a microplate reader. MDA concentrations were calculated from a standard curve generated with MDA standards and expressed relative to tissue weight.

### Glutathione measurement

Total glutathione (GSH + GSSG) levels in liver tissue were determined using the Glutathione Assay Kit (Cayman Chemical, cat# 703002) according to the manufacturer’s instructions. Briefly, ∼50–100 mg of liver tissue was rinsed with PBS, homogenized in ice-cold MES buffer (50 mM, pH 6–7, containing 1 mM EDTA), and centrifuged at 10,000 × g for 15 min at 4 °C. The resulting supernatant was deproteinated with metaphosphoric acid, neutralized with triethanolamine, and used for analysis. Samples (50 µl) were loaded in duplicate on a 96-well plate together with GSSG standards, and 150 µl of freshly prepared assay cocktail containing DTNB, glutathione reductase, and NADPH-generating cofactors was added to each well. The formation of 5-thio-2-nitrobenzoic acid (TNB) was monitored at 405 nm using a microplate reader, and glutathione concentrations were calculated from the standard curve and normalized to tissue weight.

### Statistical analysis

All data are presented as mean ± s.e.m.. Statistical significance was determined using unpaired two-tailed Student’s t-test or one-way ANOVA with Tukey’s post hoc test, as appropriate. A p-value < 0.05 was considered statistically significant. GraphPad Prism 9 (GraphPad Software) was used for all analyses and figure generation.

## Data availability

The bulk RNA-Seq data and mitochondrial proteomic data generated in this study will be deposited to the ProteomeXchange Consortium via the PRIDE partner repository and made accessible upon the publication of this study. All other data needed to reproduce the work present in this study are available in the manuscript and supplementary information. Source data are provided with this paper.

## Acknowledgements

This work was supported by grants from the National Institutes of Health (NIH-R01DK136722, F.Y.), the University of Florida (UF) Older Americans Independence Center (OAIC) Pepper Junior Scholar Award (F.Y.) with parent grant NIH-P30AG028740 (Manini, Todd Matthew) and the UF Research Startup funds. This work was also supported by the NIH Shared Instrumentation Grant NIH S10 OD021758-01A1 and NIH S10 OD030250-01A1 to UF Mass Spectrometry Research and Education Center. We are grateful to UF ICBR Electron Microscopy Core Facility for TEM analysis and other Yue lab members for their discussion.

## Author contributions

F.Y. developed the concept for the studies. Y.W., L.C., and F.Y designed the experiments. Y.W., L.C., Y.Z., Z.J., K.K., K.H., M.A.G-M., J.R., Y.L., K.B.B., T.E.R., and F.Y. performed the experiments and analyzed the data. Z.C. and K.A.E provided the key resources. Y.W. and F.Y. prepared the figures and wrote the manuscript.

