## Supplemental Figures for "Hepatocyte FAM210A deficiency disrupts mitochondrial function and triggers juvenile steatosis with compensatory repair in adulthood"

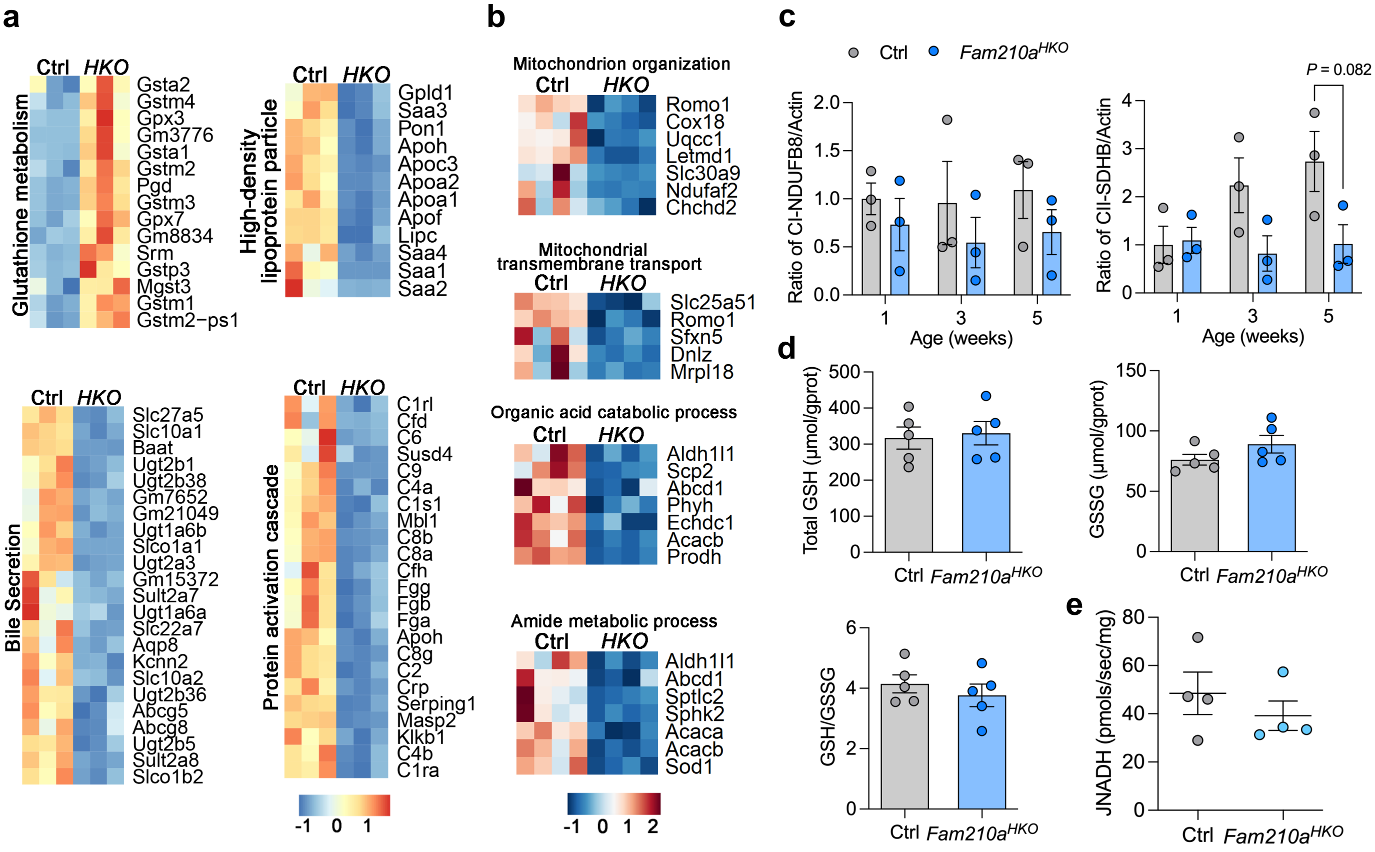


**Fig. S1 | Transcriptomics and proteomics identify enrichment of downregulated pathways and GSH activity in *Fam210a*-deficient liver. a** Heatmaps showing genes related to glutathione metabolism, high-density lipoprotein particle, bile secretion, and protein activation cascade in Ctrl and *Fam210a*$^{HKO}$ livers at 5 weeks old. **b** Heatmaps showing proteins associated with mitochondrial organization and transmembrane transport, organic and catabolic process, and amide metabolic process. **c** Quantification of relative expression of CI-NDUFB8A and CII-SDHB in Ctrl and *Fam210a*$^{HKO}$ livers at 1, 3, and 5 weeks old (n = 3; mean ± s.e.m.; two-tailed, unpaired Student's t-test). **d** Total GSH and GSSG content, and GSH/GSSG ratio in Ctrl and *Fam210a*$^{HKO}$ livers at 3 weeks old (n = 5; mean ± s.e.m.; two-tailed, unpaired Student's t-test). **e** Enzymatic activity of α-ketoglutarate dehydrogenase (aKGDH) in liver mitochondria (n = 4; mean ± s.e.m.; two-tailed, unpaired student's *t*-test).

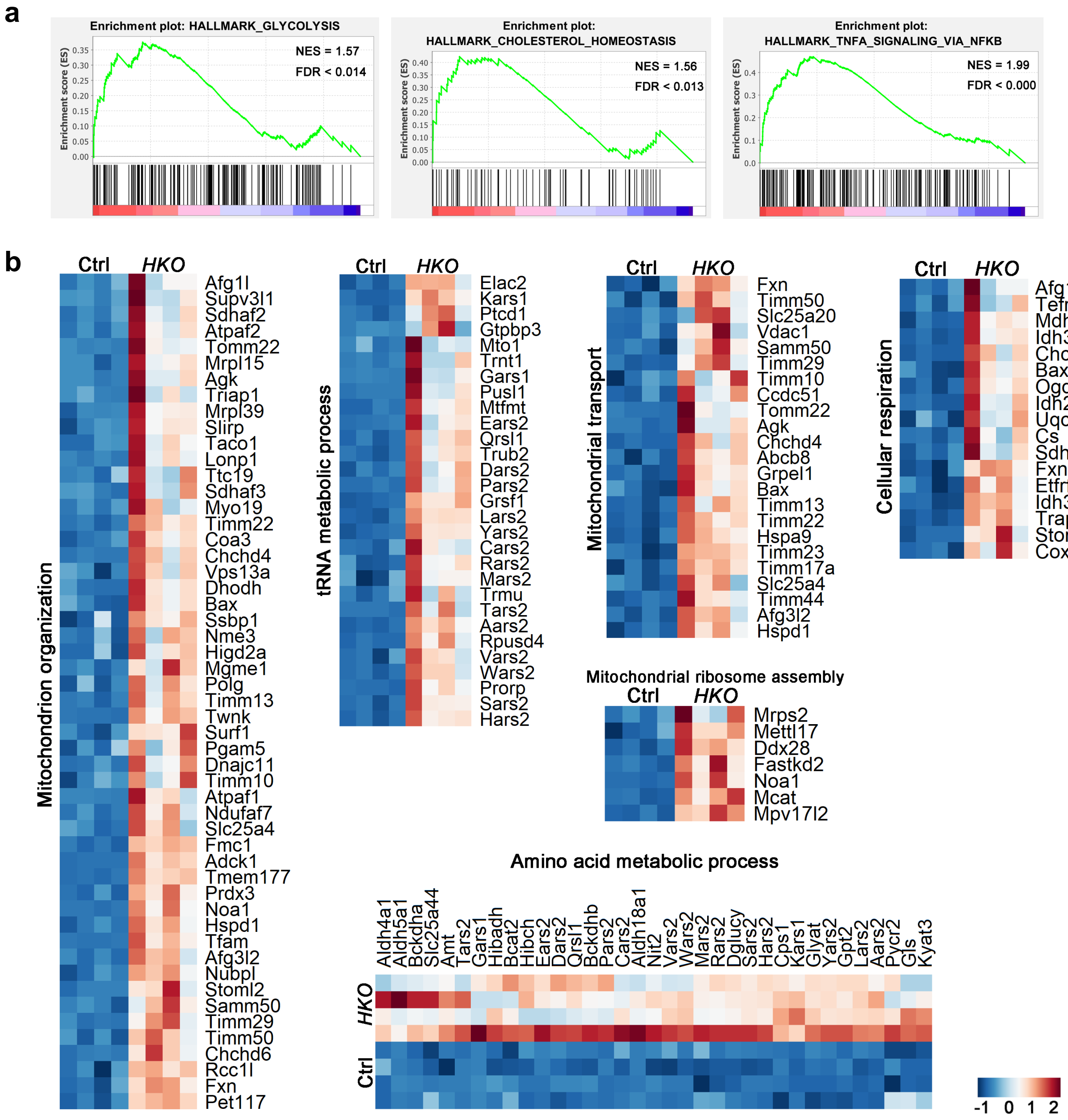


**Fig. S2 | Transcriptomics and proteomics identify enrichment of upregulated pathways in *Fam210a*-deficient liver. a** Gene Set Enrichment Analysis (GSEA) identified the enrichment of glycolysis, cholesterol homeostasis and TNFα signaling upregulated in *Fam210a*$^{HKO}$ liver at 5 weeks old (n = 3). **b** Heatmaps showing proteins related to mitochondrial organization, tRNA metabolic process, transport, ribosome assembly, cellular respiration and amino acid metabolic process upregulated in livers at 5 weeks old (n = 4).
