## Supplemental Table 1 for "Hepatocyte FAM210A deficiency disrupts mitochondrial function and triggers juvenile steatosis with compensatory repair in adulthood"

**Supplementary Table 1. Real-time PCR primer sequences.**

| **Gene name** | **Sequence (5' to 3')** |
|---|---|
| *Cdc20*: forward | TTCGTGTTCGAGAGCGATTTG |
| *Cdc20*: reverse | ACCTTGGAACTAGATTTGCCAG |
| *Plk1*: forward | CCCGCTGGCGAAAGAAATTC |
| *Plk1*: reverse | CATTTGGCGAAGCCTCCTTTA |
| *Ccnb1*: forward | AAGGTGCCTGTGTGTGAACC |
| *Ccnb1*: reverse | GTCAGCCCCATCATCTGCG |
| *Mki67*: forward | ATCATTGACCGCTCCTTTAGGT |
| *Mki67*: reverse | GCTCGCCTTGATGGTTCCT |
| *Elovl3*: forward | TTCTCACGCGGGTTAAAAATGG |
| *Elovl3*: reverse | GAGCAACAGATAGACGACCAC |
| *Slco1a1*: forward | GTGCATACCTAGCCAAATCACT |
| *Slco1a1* | CCAGGCCCATAACCACACATC |
| *Cyp4a12a*: forward | CCTCTAATGGCTGCAAGGCTA |
| *Cyp4a12a*: reverse | CCAGGTGATAGAAGTCCCATCT |
| *C8a*: forward | GGGACCCCTGGAGACGAAA |
| *C8a*: reverse | GCCCACAACGACAGGCATTA |
| *C8b*: forward | CCCCACAGACCAAATGTGAAT |
| *C8b*: reverse | CCAAGATCCATAACGCCATCCA |
| *C9*: forward | GGGTGTCAATGCACAGATGC |
| *C9*: reverse | GAGGCAAGGATCACACTCTGA |
| *β -actin*: forward | GGCTGTATTCCCCTCCATCG |
| *β -actin*: reverse | CCAGTTGGTAACAATGCCATGT |
